# SUSTAINED SYSTEMIC AND NEUROINFLAMMATION IN COGNITIVE DYSFUNCTIONAL FEMALE MICE AFTER NON-SEVERE EXPERIMENTAL MALARIA

**DOI:** 10.1101/2025.05.28.656400

**Authors:** Pamela Rosa-Gonçalves, Beatriz Nogueira Siqueira-e-Silva, Maiara Nascimento Lima, Luana da Silva Chagas, Luciana Pereira de-Sousa, Luzia Fátima Gonçalves Caputo, Igor José da Silva, João Paulo Rodrigues dos Santos, Caio César de Araujo Evaristo, Mônica da Silva Nogueira, Guilherme Loureiro Werneck, Flávia Lima Ribeiro-Gomes, Paulo Renato Rivas Totino, Lilian Rose Pratt-Riccio, Tatiana Maron-Gutierrez, Roberto Farina de Almeida, Marcelo Pelajo-Machado, Claudio Alberto Serfaty, Cláudio Tadeu Daniel-Ribeiro

## Abstract

Plasmodial infection induces systemic inflammation with great potential to contribute to the development of severe illness and lethality. It is known that the hippocampus and cortex both play a pivotal role in memory processes and are affected by the neuroinflammation associated with cerebral malaria that causes long-lasting cognitive and behavioral sequelae. Since these sequelae are also associated with the non-severe form of malaria, it is important to correlate brain morphology, particularly glial cell involvement, and neuroimmune and neuroinflammatory features with memory acquisition and consolidation processes, in this clinical presentation of malaria, the most frequent worldwide. Here, we aimed to investigate cellular and molecular neuroimmune aspects of non-severe experimental malaria-associated cognitive dysfunction. Female C57BL/6 mice were infected with *Plasmodium berghei* ANKA and treated with chloroquine before any clinical signs of cerebral malaria emerged. No histopathological alteration, in hematoxylin-eosin staining, or axonal damage, in Bielschowsky’s silver-impregnated brain sections, was observed. However, morphological alterations in GFAP^+^ and Iba-1^+^ cells suggest that: i) astrocytes in the hippocampal dentate gyrus and the *cornu Ammonis* 1 regions and ii) microglia in the *cornu Ammonis* 1 region are responding to infection. Curiously, the effect persisted only in Iba-1^+^ cells up to 22 days post-infection. An increase in pro-inflammatory cytokines levels and expression was observed, in both the prefrontal cortex and the hippocampus. Also, serum and spleen cytokine levels were increased at 4 days post-infection. At 22 days post-infection, infected and treated mice showed an increase in serum cytokine levels that had homeostatic levels at 155 days post-infection. This dynamic points to both an immune stimulus persistence and a cytokine autoregulation post-infection. Infected mice exhibited acquisition and consolidation memory deficits in behavioral tests early after treatment (22 days post-infection). In conclusion, in a context of sustained systemic inflammation, mild neuroinflammatory alterations of glial cells may be involved in cognitive sequelae following a single episode of non-severe experimental malaria.

## INTRODUCTION

The WHO (2024) estimates that more than 260 million malaria cases and around 600,000 malaria deaths have occurred in 83 countries in 2023. Cerebral malaria (CM) is the deadliest complication of the disease, and the survivors, even with access to the best treatments available, frequently develop neurological and/or neurocognitive impairments including cognitive decline (Varo *et al*., 2018; Rosa-Gonçalves *et al*., 2022a; de Sousa *et al*., 2023). Long-lasting cognitive and behavioral damage has been associated with both severe and non-severe malaria (nSM) (Rosa-Gonçalves *et al*., 2022a; Ssemata *et al*., 2023), by far, the predominant clinical form of the disease worldwide.

Plasmodial infection induces a systemic inflammation that can contribute to the development of severe forms of malaria (*P. falciparum* parasitemia and at least one of the severe disease criteria: prostration, jaundice, abnormal bleeding, hypoglycemia, acidosis, hyperlactatemia, severe anemia, impaired consciousness, respiratory distress, multiple convulsions, renal impairment, pulmonary edema, shock or hyperparasitemia) (Gazzinelli *et al*., 2014; Coban *et al*., 2018; Conroy *et al*., 2019). Inflammatory cytokines increase the expression of ligands such as intercellular adhesion molecule 1 (ICAM-1) and others on the surface of endothelial cells to which the parasitized red blood cells (pRBC) adhere via *P. falciparum* erythrocyte membrane protein 1 (*Pf*EMP-1) expressed on the knobs protuberance present on the pRBC (Gazzinelli *et al*., 2014; Watermeyer *et al*., 2016; Coban *et al*., 2018). Interactions between the parasite’s antigens and host receptors result in pRBC sequestration in the capillaries. When maturing, parasites release bioactive molecules, such as hemozoin, impairing the expression of MHC class II, and triggering a Th1 profile immune response with a high load of inflammatory cytokines. This cytokine release, whether into the peripheral circulation or locally in organs, results in symptomatic disease that can take severe forms, such as CM and kidney and lung damage (Schwarzer *et al*., 1992; Lewis *et al*., 2019).

Our group described long-lasting cognitive and behavioral sequelae after non-severe experimental malaria (nSEM) by applying the broadly studied model of CM (*Plasmodium berghei* infected C57BL/6 mice) but treating infected mice at the fourth-day post-infection, just before any neurological sign of CM syndrome appears, thus adapting the CM model for nSM studies (de Sousa *et al*., 2018; Rosa-Gonçalves *et al*., 2022b).

Regarding cognitive processes, both hippocampus and cortex play a pivotal role in memory formation (Izquierdo & Medina, 1997), and are affected by neuroinflammation processes identified in CM (de Miranda *et al*., 2017), particularly the frontal and temporal lobes (Fernando *et al*., 2010). However, no effective treatment is presently available for malaria-associated cognitive sequelae, and the knowledge about its neuropathogenesis is still poorly understood, since *Plasmodium* parasites infect red blood cells but not neural cells. Therefore, the understanding of astrocytes and microglial cell activation and neuroimmune signaling in memory acquisition and consolidation processes during nSM is mandatory. In this study, we investigated cellular and molecular neuroimmune aspects of nSM cognitive dysfunction. We carried out histopathological, immunostaining, cellular and molecular analyses of cognitive brain regions along with behavioral tests and cytokine analyses in nSEM.

## METHODS

### Host and parasite strain

Seven-week-old, 18-20g, female C57BL/6 mice (n=125) were bred and maintained in the animal facility of the *Instituto de Ciência e Tecnologia em Biomodelos* (*Fiocruz*) and *Centro de Experimentação Animal* (*Instituto Oswaldo Cruz, IOC*, *Fiocruz*) under the license’s numbers L-010/2015 and L-004/2020. Female mice were chosen based on previous results of malaria-associated cognitive and behavioral impairment (de Sousa *et al*., 2018, 2021; Rosa-Gonçalves *et al*., 2022b). All experiments followed ethical and animal welfare practices and were approved by the Ethics Committee of the *IOC* (*CEUA*, *Fiocruz*). This study was carried out in compliance with the ARRIVE guidelines. Animals were maintained in polypropylene cages, five per cage, with one Igloo^TM^, kept in racks with air filtration system, chow and water *ad libitum*, 50% (±10) humidity, 20° C (±2) and 12-hour light/dark cycle in the animal facility of *IOC*, *Fiocruz*. The experimental infection was performed with the ANKA strain of *Plasmodium berghei* (*Pb*A), that expresses a green fluorescent protein (GFP) from the Malaria Research and Reference Reagent Resource Center (MR4 number: MRA-865)-BEI Resources.

### Infection and antimalarial treatment

C57BL/6 mice were randomly assigned into two experimental groups exposed to an intraperitoneal injection of either 10^6^ *Pb*A-GFP pRBC in phosphate-buffered saline (PBS), for the infected group (INF), or vehicle PBS, for the control group (CTRL). The pRBC were obtained from the blood of “passage” mice (n=4), as previously described (Rosa-Gonçalves *et al*., 2022). Although this model is highly susceptible to CM, the treatment with chloroquine (CQ, Farmanguinhos-Fiocruz), 25 mg/Kg/day in 200 μL of PBS, from the fourth-day post-infection (dpi) on, for seven days, prevents the development of the fatal cerebral syndrome (de Sousa *et al*., 2018). Both the INF and CTRL groups received CQ treatment by gavage. This treatment protocol did not affect the mice’s behavior in cognitive tests (de Sousa *et al*., 2018; Rosa-Gonçalves *et al*., 2022b). Parasitemia was measured at 4, 10, 22 and 155 dpi by flow cytometry or microscopy of blood on slides (Rosa-Gonçalves *et al*., 2022b). Blood was collected (3 μL) by a minimal section of mouse tails and was diluted in PBS (50 μL) and maintained at 4°C for up to a week for later acquisition on the cytometer. Before acquisition in a Cytoflex S flow cytometer (Beckman Coulter, Pasadena, CA, USA), 5 μL of homogenized content was diluted in 500 μL of PBS to obtain the percentage of GFP^+^ red blood cells. Control mice were subjected to the same procedure of minimal tail sectioning.

### Experimental design and procedure

CTRL and INF groups were treated with CQ from the fourth dpi, or after PBS injection, to avoid the development of the severe cerebral syndrome, thus constituting a nSEM (**Figure 1**) (de Sousa *et al*., 2018). Twelve days later, behavioral protocols were performed to assess cognitive parameters. The cellular and molecular aspects of neuroimmune response were assessed in the brain (prefrontal cortex and hippocampus), spleen, and serum obtained by cardiac puncture in animals from both cohorts at: i) 4 dpi (CTRL and INF); ii) 22 dpi (CQ-treated) and iii) 155 dpi (CQ-treated). All CTRL groups were mock-infected, treated with CQ whenever applicable, and treated in the same manner as the INF groups.

**Figure 1.**
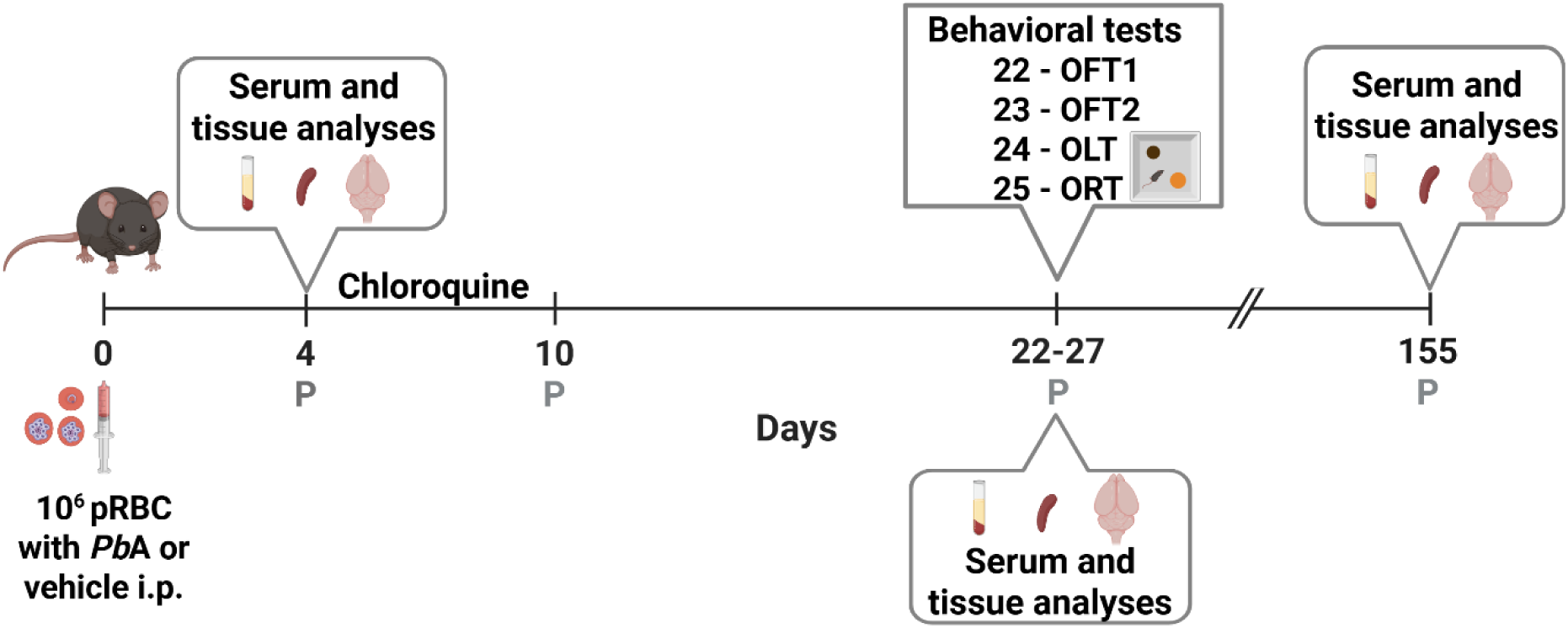
Experimental design: study timeline of cellular and molecular neuroimmune aspects of non-severe experimental malaria-associated cognitive dysfunction. C57BL/6 mice (n=118) were inoculated intraperitoneally (i.p.) with 10⁶ *Plasmodium berghei* ANKA-parasitized red blood cells (pRBC) or vehicle (PBS), and treated with chloroquine (CQ) from days 4 to 10 post-infection (D4-D10). Behavioral tests, such as open field (OFT), object location (OLT) and object recognition (ORT), were performed on day 22 post-infection in one cohort (CTRL and INF, both CQ-treated, n=48), followed by serum and tissue collection for cellular and molecular analyses. Other cohorts were analyzed on day 4 (n=60) and 155 (n=10) post-infection for early- and late-stage immune profiling. Parasitemia (P) was assessed on days 4, 10, 22, and 155 using flow cytometry or microscopy. Figure created in https://BioRender.com.

From the animals assigned to cohort i (n=60), the following groups were formed and considered to: CTRL (n=30), INF (n=30). From the animals assigned to cohort ii (n=48), the following groups were formed and considered: CTRL-CQ (n=24), INF-CQ (n=24). From the animals assigned to cohort iii (n=10), the following groups were formed and considered: CTRL-CQ (n=5), INF-CQ (n=5). Two mice had low levels of parasitemia and were excluded from the analyses.

### Behavioral analysis

The sequence of behavioral tests assessing memory acquisition and memory consolidation was conducted at 22 dpi, following the methodology reported by Rosa-Gonçalves *et al*. (2022b) and adapted from Vogel-Ciernia & Wood (2014) (**Figure 2**). The tests were carried out in the afternoon, under a 60 lux luminance of incandescent lamps and after a 2h period of acclimatization with access to *ad libitum* chow and water. The apparatus was cleaned with 70% alcohol and dried between each animal test session. The tests were recorded, and the automated acquisitions and analyses were performed using AnyMaze® 5.1 software (Stoelting Co., Wood Dale, IL). For the OLT and ORT, manual analyses were performed by a researcher blinded to group assignment. The time spent snout facing the object at a distance ≤ 2 cm and/or touching the object with the snout or forepaws was manually measured by a researcher blinded to treatment, using a digital stopwatch, by observing video recordings of the sessions. The exploration time in each object and its percentage were calculated by dividing the exploration time of the novel or familiar object/location by the sum of the exploration time for the two objects, multiplied by 100.

**Figure 2.**
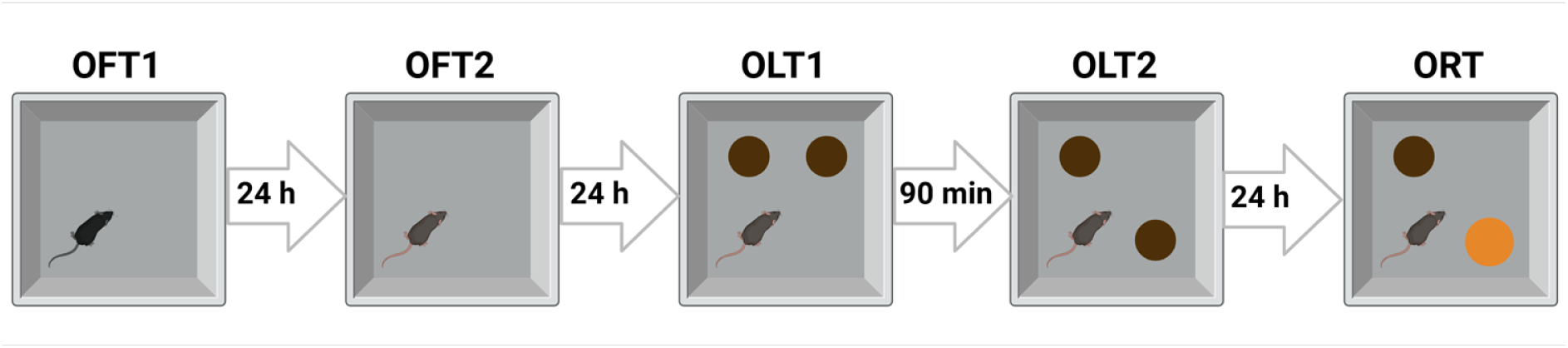
Behavioral tests sequence. At 22 dpi, mice underwent two open field test (OFT) sessions, each lasting 10 min, with a 24 h interval between sessions. In the object location test (OLT), mice explore two identical objects for 10 minutes during the training session (OLT1). After a 90 min of retention interval (RI), the location of one object is changed, and mice explore them again for 10 min (OLT2). In the object recognition test (ORT), the object at the novel location is changed to a new object and both are explored for 10 min, 24 h of RI.

#### Open field test

The OFT was carried out as already described by Rosa-Gonçalves *et al*. (2022b). Mice at 22 dpi freely explored individually the gray acrylic box arena (50 × 50 × 50 cm) for 10 min for two sessions (OFT1 and OFT2) 24 h apart. This task was performed to characterize the habituation memory by: the total time, the first three minutes and minute-a-minute distances traveled in both sessions to reduce the novelty to the environment. The parameter comparison of the same animal between sessions allows the inference of the performance in habituation memory, since the environment, already explored, is consequently less explored in OFT2.

#### Object location test

The OLT was adapted from Vogel-Ciernia & Wood (2014). This task was carried out in the same arena of OFT, 24 h apart. It was performed to assess the acquisition memory process, based on the mice’s natural preference for novelty. After 24 h of OFT2 session, mice freely explored individually two identical objects for 10 min in an OLT training session (OLT1). After 90 min, the location of one of the objects was changed to a new location and mice freely explored individually for 10 min in an OLT test session (OLT2). The object at the novel location is more explored than at the familiar location, based on the natural tendency of animals to explore the novelty. Before the test session, the mice were subjected to a training session, in which all groups performed similarly (**Figure S1**).

#### Object recognition test

The ORT was adapted by Rosa-Gonçalves *et al*. (2022). This task was carried out in the same arena of OLT, 24 h apart. It was performed to assess the consolidation memory process, based on the mice’s natural preference for novelty. After 24 h of OLT2, one of the familiar objects was replaced by a new object with different texture, shape, color and size. Mice were permitted to explore, individually, both familiar and novel objects for a period of 10 min within an ORT session. The novel object is more explored than the familiar, based on the natural tendency of animals to explore the novelty. In this protocol, the OLT2 fulfills the role of ORT training session.

It is important to inform that CQ treatment of *Pb*A-infected mice — that clears the parasitemia sustainably — did not affect the cognitive and behavioral performance of C57BL/6 mice in different paradigms (De Sousa *et al*. 2018; Rosa-Gonçalves *et al*. 2022b).

### Enzyme-Linked Immunosorbent Assay (ELISA)

Prefrontal cortices, hippocampi and spleens of mice of cohorts 4 and 22 dpi were dissected and stored at -80°C. The samples were homogenized (speed 5 of IKA model T10 Basic) in ice-cold PBS, 0.1% Triton and protease inhibitor solution (Merck, 11836153001) and centrifuged at 2000 g. The supernatants were sampled and stored at -80°C. The BCA method (Bicinchoninic Acid - Thermo Scientific, Pierce, USA) was used to normalize cytokines values to determine the total protein concentrations. ELISA DuoSet kits were used to detect the concentration of the cytokines IL-1β, TNF-α, IL-6, KC and IL-10, according to the manufacturer’s instructions (R&D Systems, Minneapolis, MN, USA) (Lima *et al*., 2020). Plates were coated with 50 μL of capture antibody and incubated overnight at 4°C. The plates were washed five times in PBS and Tween 20 (0.5%) and blocked (PBS and 1% bovine serum albumin solution) for 2h and washed again. The standard sample and those to be tested were added and incubated for 2h at room temperature. The detection antibody was added and incubated for 2h and washed. Then, the streptavidin peroxidase-conjugated solution was added and incubated for 20 min and washed. The chromogenic substrate solution (TMB, Sigma) was added, and the reaction was stopped with a sulfuric acid solution (1M). Optical densities (OD) were acquired on a microplate reader at 450 nm.

### Cytokines levels by flow cytometry

The samples of mice at 4, 22 and 155 dpi were measured with Cytometric Bead Array (CBA) Mouse Th1/Th2/Th17 (BD Biosciences, 560485) according to the manufacturer’s instructions (de Sousa *et al*., 2021). The serum was obtained from centrifugation of total blood cardiac puncture (from deeply anesthetized mice), and the right hemisphere of the hippocampus was dissected and stored at -80°C. The tissue was homogenized in 60 µL of ice-cold PBS, 0.1% Triton-X100 and 1% protease inhibitor (Thermo Fisher, 78430) solution. The supernatant sample was collected and stored at -80°C. The BCA method (Thermo Fisher, Pierce, USA) was used to construct a standard curve and determine the protein concentrations. The samples were acquired in CytoFLEX S cytometer (Beckman Coulter, Pasadena, CA, USA), and data analysis was performed using the FlowJo 10.0 software (BD Biosciences).

### Histological analysis of paraffin sections

The dissected brains of mice cohorts at 4, 22 and 155 dpi were isolated and immersed in 10% Carson’s buffered formalin for fixation and then sagittal sections were obtained. The left hemispheres were placed in histological cassettes and processed through the dehydration, clearing and impregnation using a Shandon Citadel 2000 automatic tissue processor. The tissues were embedded in paraffin and positioned on a cold plate until the blocks solidified and could be removed from the molds. The resulting blocks were sectioned using a Leica RM2125 RT microtome.

#### Hematoxylin-Eosin (HE) staining

Histological sections were routinely stained with Mayer’s hematoxylin and eosin-phloxine (Caputo *et al*., 2011), mounted with Damar’s gum and, analyzed under bright-field microscopy using an Axioskop microscope (Carl Zeiss, Germany) equipped with an Axiocam 712 camera and ZEN software. Images were analyzed using 10x, 40x and 100x objective lenses.

#### Silver impregnation

Paraffin-embedded brain sections were impregnated with silver using Bielschowsky’s method, as previously reported (Prophet *et al*., 1994), with minor modifications. This technique allows the analysis of neuronal fibers and synapses during infection. Brain tissue sections were deparaffinized and hydrated. Seven µm sections were treated in silver nitrate solution 20% and incubated at 56°C for one hour or until the tissue turned amber in color. They were immersed in distilled water and after in ammoniacal water (containing concentrated ammonium hydroxide), then in ammoniacal silver (silver nitrate used in sensitization and ammonium hydroxide drop by drop until the solution became clear). Before adding the slides, the revealing solution was added (formaldehyde, distilled water, nitric acid and citric acid). They were toned with 1% gold chloride and then immersed in a 5% sodium thiosulphate solution. The slides were then dehydrated, clarified, mounted and finally analyzed under an optical microscope (Axioskop, Carl Zeiss, Germany). Images were analyzed using 10x, 40x and 100x objective lenses.

### Immunofluorescence

Mice at 4 and 22 dpi were deeply anesthetized with an overdose of ketamine and xylazine i.p. and then perfused transcardially with NaCl (0.9%) containing heparin (0.01%), for 2 min, followed by buffered 4% paraformaldehyde in saline-phosphate buffer (0.1 M), pH 7.4 for 10 min, performed with a peristaltic pump (Bonther, BP600). Brains were dissected and subsequently cryoprotected in 20% sucrose for 24h at 4°C, frozen in dry ice and stored at −80 °C. Coronal sections of 10 μm were cut using a cryostat (CM1860, Leica) and mounted onto Poly-L-Lysine (Sigma-Aldrich, P8920) coated slides. The brain regions were identified using Allen Reference Atlas – Mouse Brain (atlas.brain-map.org) (Allen Institute for Brain Science, 2004). The immunofluorescence protocol was based on previous detailed descriptions (Chagas *et al*., 2019). All tissue sections were washed in PBS, pH 7.4, permeabilized in PBS with Triton X-100 (Sigma, T8787, PBS-T) 0.5%, then blocked in 10% normal goat serum (1000C, Thermo, NGS), bovine serum albumin 0.3% and PBS-T for two hours at room temperature. The sections were incubated overnight with the appropriate primary antibodies: either rat anti-GFAP (1:1000, Invitrogen, 130300) or rabbit anti-Iba (E4O4W, Alexa Fluor® 488 Conjugate, 1:100, Cell Signaling, #20825) in PBS-T at 4 °C. After three washes in PBS, the sections previously incubated with unconjugated antibodies were incubated with secondary Alexa Fluor® 594 conjugated goat anti-rat (1:200, Invitrogen, A48264) for two hours at room temperature. All sections were washed three times in PBS and counterstained with DAPI 1µg/mL (4’,6-diamidino-2-phenylindole dihydrochloride, Sigma, D9542) to visualize the nuclei, and coverslips were mounted in n-propyl-gallate (Sigma, P3130) and glycerol (Sigma, G5516) medium. Images were acquired using a deconvolution microscope system (DM5500B, Leica) under 400x magnification.

#### Quantification of immunofluorescence images

Densitometric analysis of the staining level was performed on 8-bit images using ImageJ 1.54p software (http://imagej.org). The integrated density was calculated as the sum of the pixel values in an image, setting the threshold to Auto in Default mode for GFAP and Yen mode for Iba-1 analysis per image. We obtained pictures from different adjacent sections in the same region and then calculated the mean densitometric values. The results were expressed by dividing the average of the integrated density values per field by the average of the respective control group. The morphology quantification was performed using the software ImageJ plugin AnalyzeSkeleton (2D/3D), according to Young & Morrison (2018) protocol.

### Gene expression by RT-qPCR

Prefrontal cortices and hippocampi of mice of cohort at 4 dpi and spleens of cohorts at 4 and 155 dpi were dissected and stored in RNAlater (Thermo Fisher) at -20°C. The tissues were homogenized and then the messenger RNA (mRNA) was isolated and purified using silica columns provided by the PureLink RNA mini Kit (Ambion), following the manufacturer’s instructions. The mRNA was converted into complementary DNA (cDNA) by reverse transcription (RT-PCR) following the instructions of the High Capacity cDNA Reverse Transcription kit (Applied Biosystems™), and subsequently, the cDNA was quantified using the Qubit ssDNA Assay (ThermoFisher) and normalized to 100 ng/mL. The evaluation of the gene expression of the T-cell receptor Vβ8. 1 (ARXGU7F) and the cytokines IL-1β (Mm00434228_m1), IL-6 (Mm01210732_g1), IL-10 (Mm01288386_m1), TNF-α (Mm00443258_m1) and CXCL1 (Mm04207460_m1) was carried out by real-time PCR using the TaqMan Gene Expression Assays system (ThermoFisher), containing specific primers and probes (FAM) for the genes of interest. The constitutive gene GAPDH (Mm99999915_g1) was used as endogenous control to normalize amplification and correct for variations and a negative control without DNA was included in all reactions. The reaction was carried out on a 7500 real-time PCR thermal cycler (Applied Biosystems). The data generated was analyzed using the Δ-ΔCT comparative calculation method to measure relative quantification with the aid of the TaqMan Expression Suite software.

### Statistical analysis

Behavioral results were expressed as measures of mean ± standard errors of the mean. Molecular analyses were expressed in heatmaps. The other results are expressed as measures of mean ± standard deviations and violin plots with median and quartiles. One-way analysis of variance (ANOVA) followed by Bonferroni’s multiple comparison post hoc test was used for comparisons of means between more than two groups (immunofluorescence analyses), while two-way analysis of variance followed by Bonferroni’s multiple comparison post hoc test was used to compare the distances traveled between groups and OFT sessions (repeated measures) and exploration time (repeated measures) in OLT and ORT. The Mann-Whitney test was used to compare the medians between two groups (cytokine and chemokine analyses by ELISA, RT qPCR flow cytometry and histological analyses). Outliers were removed based on Grubb’s test. Differences with p values <0.05 were considered statistically significant. Graphs and statistical analysis were carried out using the GraphPad Prism 9.3.0 (La Jolla, CA, USA).

## RESULTS

Similarly to data from previous literature, the plasmodial infection is reverted by chloroquine treatment, validating the nSEM. The infected mice showed around 2.5% of parasitemia at the fourth dpi (**Figure 3**) (de Sousa *et al*. 2018, 2021; Rosa-Gonçalves *et al*. 2022b). Parasitemia was negative post-CQ treatment before (D10) and after the behavioral tests (D22 and D155) in all mice.

**Figure 3.**
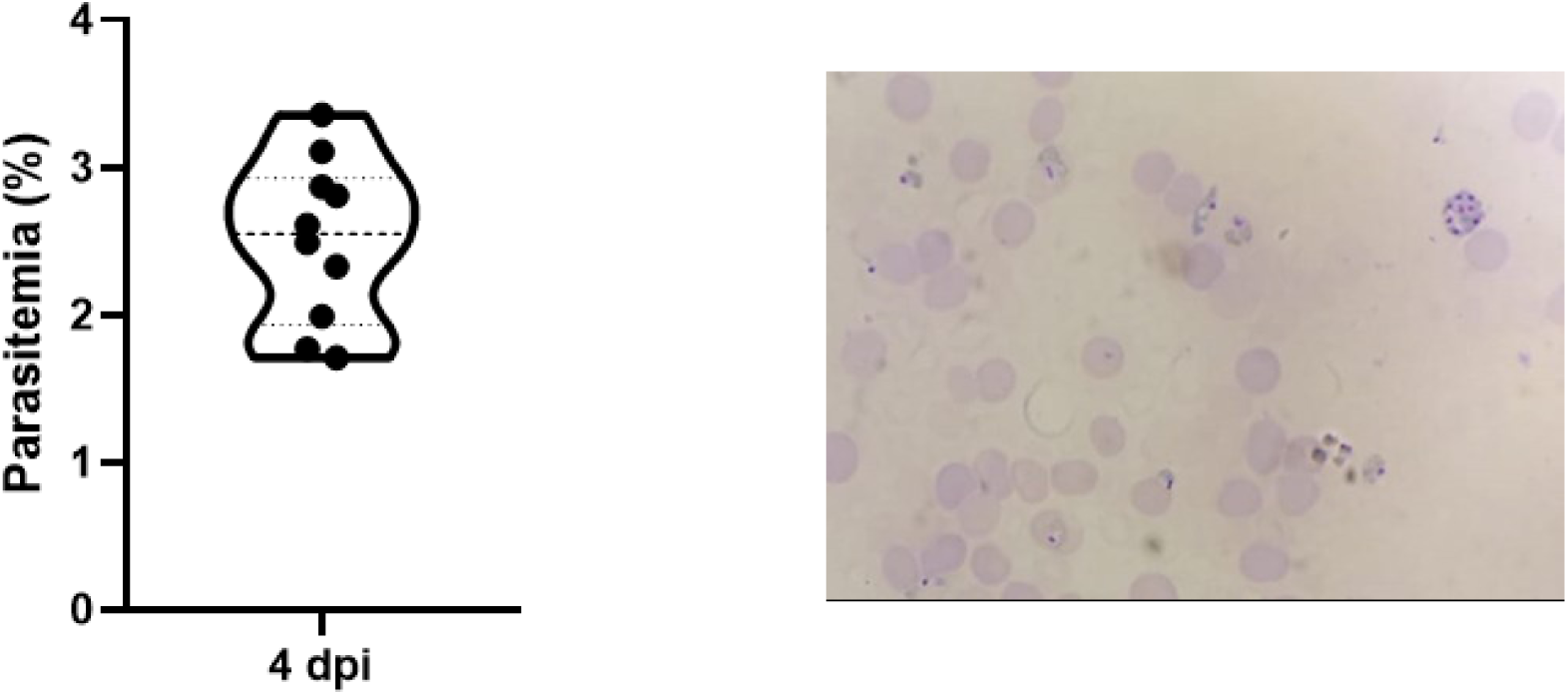
Parasitemia in the non-severe experimental malaria model. Parasitemia levels at 4 days post-infection assessed by flow cytometry (A), and representative photograph of parasitized red blood cells in blood smear microscopy (n=10). Data points are identified as individuals’ values. The violin plot shows the median and quartiles in dashed lines and a frequency distribution. Magnification: 1000X.

### Non-severe plasmodial infection impairs memory acquisition and consolidation

To evaluate the effect of non-severe *Pb*A infection on processes of memory formation, uninfected and infected / treated mice were subjected to behavioral tests as early as around two weeks after CQ treatment (at 22 dpi).

Non-severe *Pb*A infection did not alter the total distance traveled in the Open Field when compared to the uninfected control group. There was, therefore, no effect of infection on long-term habituation memory (**Figure 4A**). **Figure 4B** illustrates distance traveled during the first three minutes in the Open Field test, highlighting a significant reduction between OFT1 and OFT2 distances traveled observed both in uninfected / treated (CTRL-CQ) and infected / treated (INF-CQ) mice, indicative of habituation to a novel environment.

**Figure 4.**
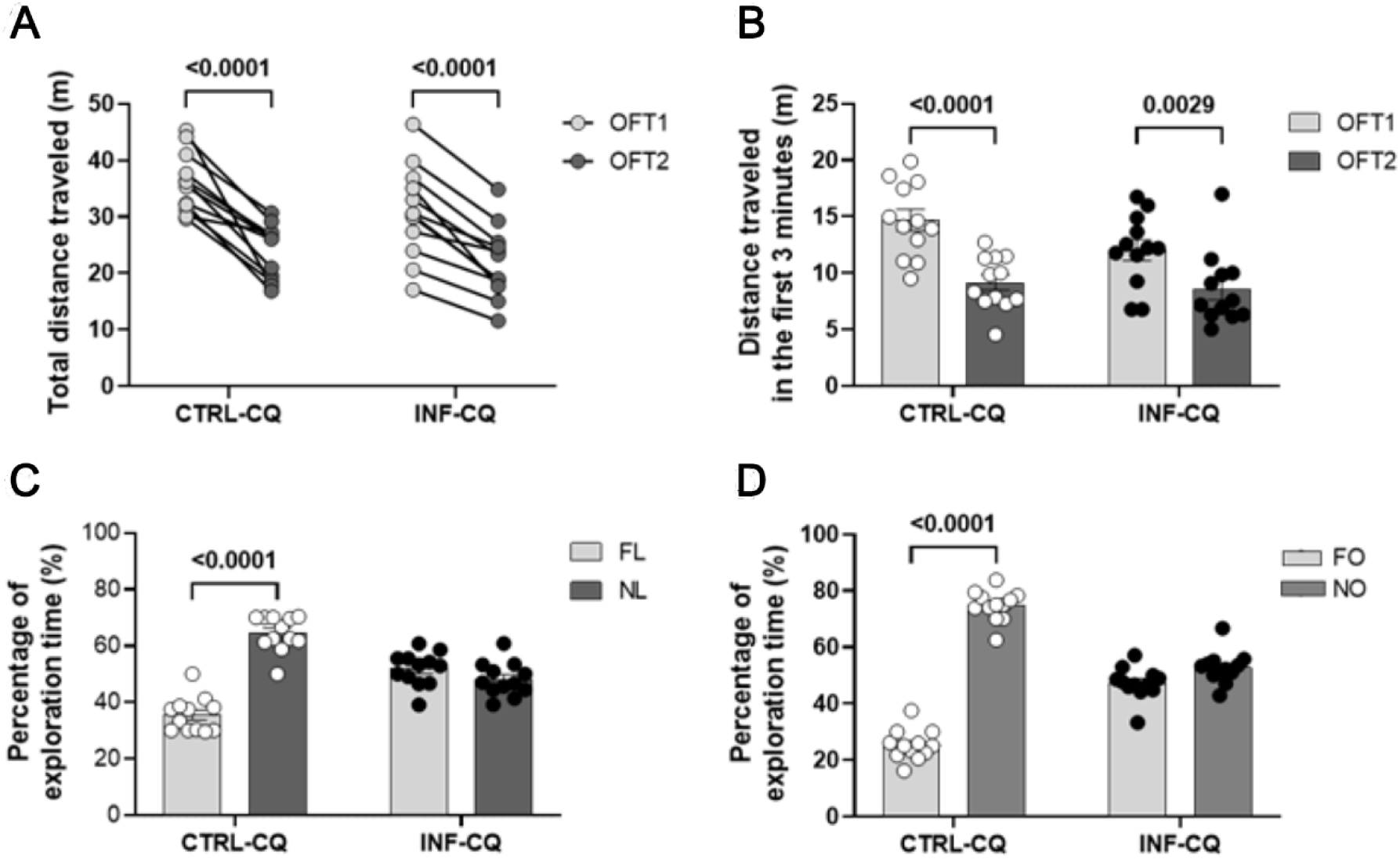
Effect of non-severe *Plasmodium berghei* ANKA infection on short- and long-term memories in mice. (**A**) Total distance traveled (m) in the Open Field test in the training (OFT1) and test (OFT2) sessions. (**B**) Distance traveled (m) in the first three minutes in the Open Field test (OFT1 and OFT2). (**C**) Percentage of time spent exploring objects in the test session of the Object Location Test. (**D**) Percentage of time spent exploring objects in the test session of the Object Recognition test. CTRL-CQ: uninfected and CQ-treated mice; INF-CQ: infected and CQ-treated mice; FL: familiar location; NL: novel location; FO: familiar object; NO: novel object. Data points are identified as individuals’ values. Columns represent mean ± standard error. Two-way RM ANOVA/Bonferroni was used for intra-group comparisons of the different sessions and objects. Two-way ANOVA was used for comparisons between CTRL-CQ and INF-CQ in OFT1 and OFT2, and the p-values were non-significant (n=12). p-values <0.05 in the graphs. Representative of two experiments.

Ninety minutes after the training session, INF-CQ mice did not show any preference of exploration between the objects and were, therefore, unable to distinguish the object in the novel location (NL) from the one in the familiar location (FL). The CTRL-CQ mice explored the object in the NL more than in the FL, as expected (**Figure 4C**). *Pb*A-infected and CQ-treated (INF-CQ) mice showed, therefore, an impairment in the short-term memory acquisition, detected in the Object Location test following infection / treatment (**Figure 4C**).

In the Object Recognition test, the CTRL-CQ mice demonstrated, as expected, an intact recognition memory, exploring more the novel object (NO) than the familiar object (FO), showing a greater preference over FO (**Figure 4D**). No differences were observed in the exploration of objects by the INF-CQ group, clearly indicating an affected recognition memory.

### Plasmodial infection is characterized by sustained systemic inflammation

Increases in splenic inflammatory cytokines (TNF-α, IL-6, IL-1β, IL-10) and chemokine (KC) levels were observed following the acute infection (cohort i at 4 dpi) (**Figure 5A**). After antimalarial treatment (cohort ii at 22 dpi), cytokines and chemokine levels remained at homeostatic control levels (**Figure 5A**). Gene expression of cytokines (IL-6, IL-10, IL-1β) and T cell receptor (Vβ8.1) was increased in the spleen at 4 dpi, suggesting systemic inflammation (**Figure 5B**). At long-term (cohort iii at 155 dpi), gene expression is similar to homeostatic levels (**Figure 5B**). Infected animals showed a darker spleen pigmentation, typical of hemozoin, at days 4 and 22 dpi and even in a long-term (155) post-infection (**Figure 5C**).

**Figure 5.**
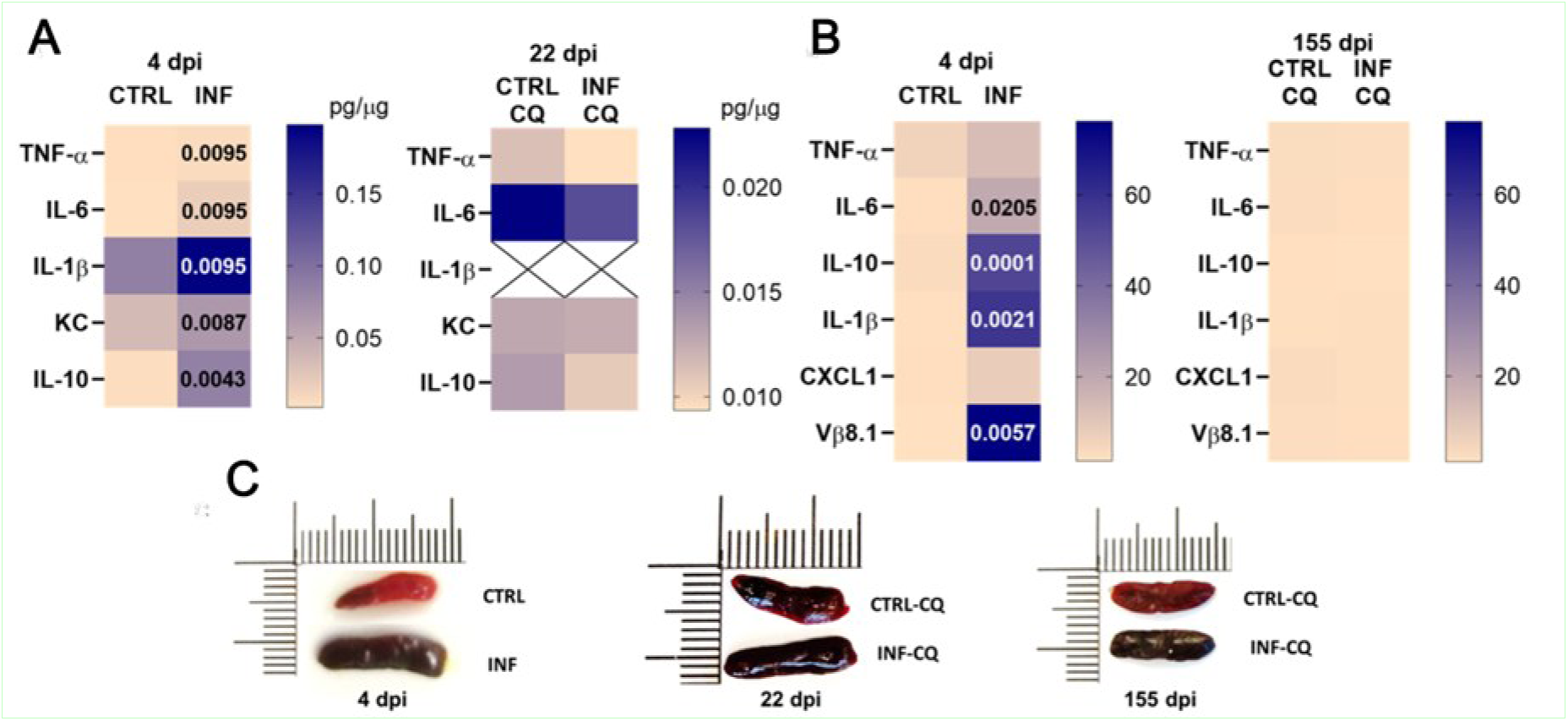
Effect of non-severe experimental plasmodial infection on spleen. (**A**) Heatmap representing the effect of non-severe *Plasmodium berghei* ANKA infection in mice on spleen cytokine levels at 4 days post-infection (n=4-6), before treatment (TNF-α, IL-6, IL-1β, KC and IL-10) and 22 dpi (n=6-10) by ELISA. (**B**) Gene expression of TNF-α, IL-6, IL-10, IL-1β, CXCL1 and TCR Vβ8.1, at 4 (n=8-10) and 155 days post-infection (n=4-5) on spleen by RT-qPCR. (**C**) Representative photograph of spleens. Infected mice exhibited a darker pigmentation typical of hemozoin, even after 22 and 155 days post-infection. CTRL: uninfected; INF: infected; CTRL-CQ: uninfected and treated with CQ; INF-CQ: infected and treated with CQ. Mann-Whitney test. p-values <0.05 in the heatmaps.

To characterize the systemic inflammatory profile, serum Th1, Th2, Th17 and regulatory cytokine levels were measured at the peak of infection (4 dpi) and after CQ treatment in the cohorts analyzed at 22 and 155 dpi. Increased serum levels of TNF-α, IFN-ɣ, IL-6, IL-2, and IL-10 and a reduction in TGF-β1 levels were detected in the infected group 4 dpi, before treatment, as compared to the uninfected mice (**Figure 6**). No variation in IL-17A and IL-4 levels was identified.

**Figure 6.**
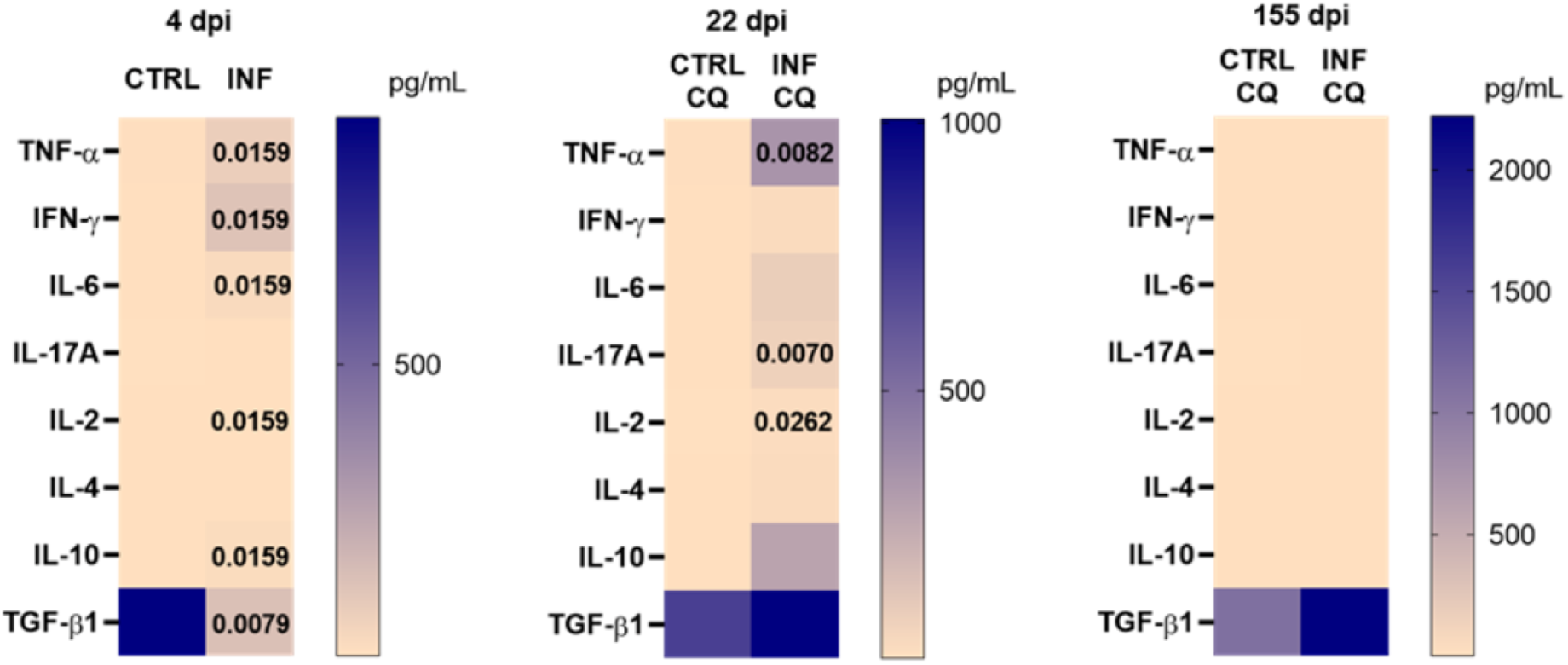
Effect of non-severe experimental plasmodial infection on serum cytokine levels. Heatmap representing the effect of non-severe *Plasmodium berghei* ANKA infection in mice on peripheral cytokine levels (TNF-α, IFN-ɣ, IL-6, IL-2, IL-4, IL-17A, IL-10, and TGF-β1) at 4-(n=4-5), 22-(n=6-7) and 155-days post-infection (n=3-5) in serum by flow cytometry (4 days post-infection values were obtained before treatment). As heatmap staining shows pg/mL concentrations, some cytokines with lower levels of expression do not exhibit pronounced staining differences as observed in higher levels of analytes, even in significant differences. CTRL: uninfected; INF: infected; CTRL-CQ: uninfected and treated with CQ; INF-CQ: infected and treated with CQ. Mann-Whitney test. p-values <0.05 in the heatmaps.

After CQ treatment, at 22 dpi, TGF-β1 levels in the infected mice did not differ from values of control mice. An increase was, however, registered in TNF-α, IL-17A and IL-2 levels. Late after antimalarial treatment (at 155 dpi), the levels of TGF-β1, IL-17A, IL-6, and IL-2 were comparable to the homeostatic ones (**Figure 6**).

The TNF-α/IL-10 and IFN/IL-10 ratio levels showed a prominent Th1-type pro-inflammatory serum cytokine profile on the fourth dpi that was not maintained after CQ treatment (**Figure 7**).

**Figure 7.**
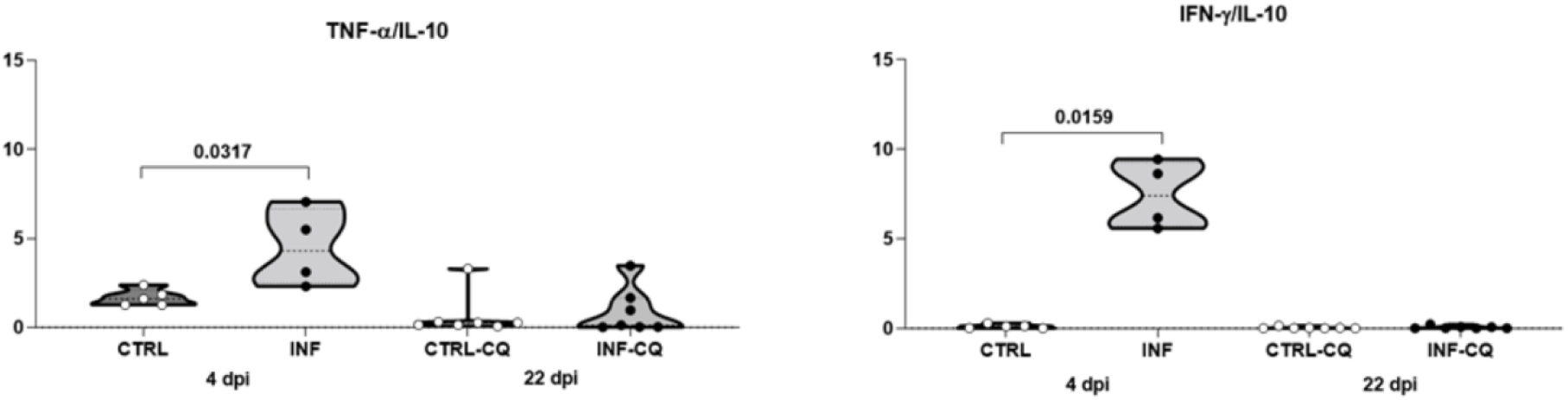
Effect of non-severe *Plasmodium berghei* ANKA infection on pro- and anti-inflammatory response balance. Ratio between the levels of peripheral cytokines TNF-α/IL-10 and IFN-ɣ/IL-10 from the cohorts analyzed at 4-(n=4-5) and 22-(n=6-7) in serum by flow cytometry. Ratio for day 155 was not calculated since no cytokine level alteration was recorded at this moment. CTRL: uninfected; INF: infected; CTRL-CQ: uninfected and treated with CQ; INF-CQ: infected and treated with CQ. Data points are identified as individuals’ values. The violin plots show the median and quartiles in dashed lines and a frequency distribution. Mann-Whitney test. p-values <0.05 in the graphs.

### No morphological change or axonal damage in the brain of mice with non-severe malaria, as assessed by hematoxylin-eosin and Bielschowsky staining

Illustrative images of the general aspect of the regions examined are shown at low magnification, but no injury was detected at the higher magnification exam. In hematoxylin-eosin (HE) staining in the brain, no evidence of inflammatory infiltrates, edema, hemorrhages, vascular congestion, amorphous material deposits, degenerative areas or necrosis was observed. The brain tissue showed neurons with a regular appearance, well-defined basophilic cytoplasm, triangular shape, large vesicular nucleus, with dispersed chromatin and prominent nucleolus, with no structural alterations between infected and control mice (**Figure 8**).

**Figure 8.**
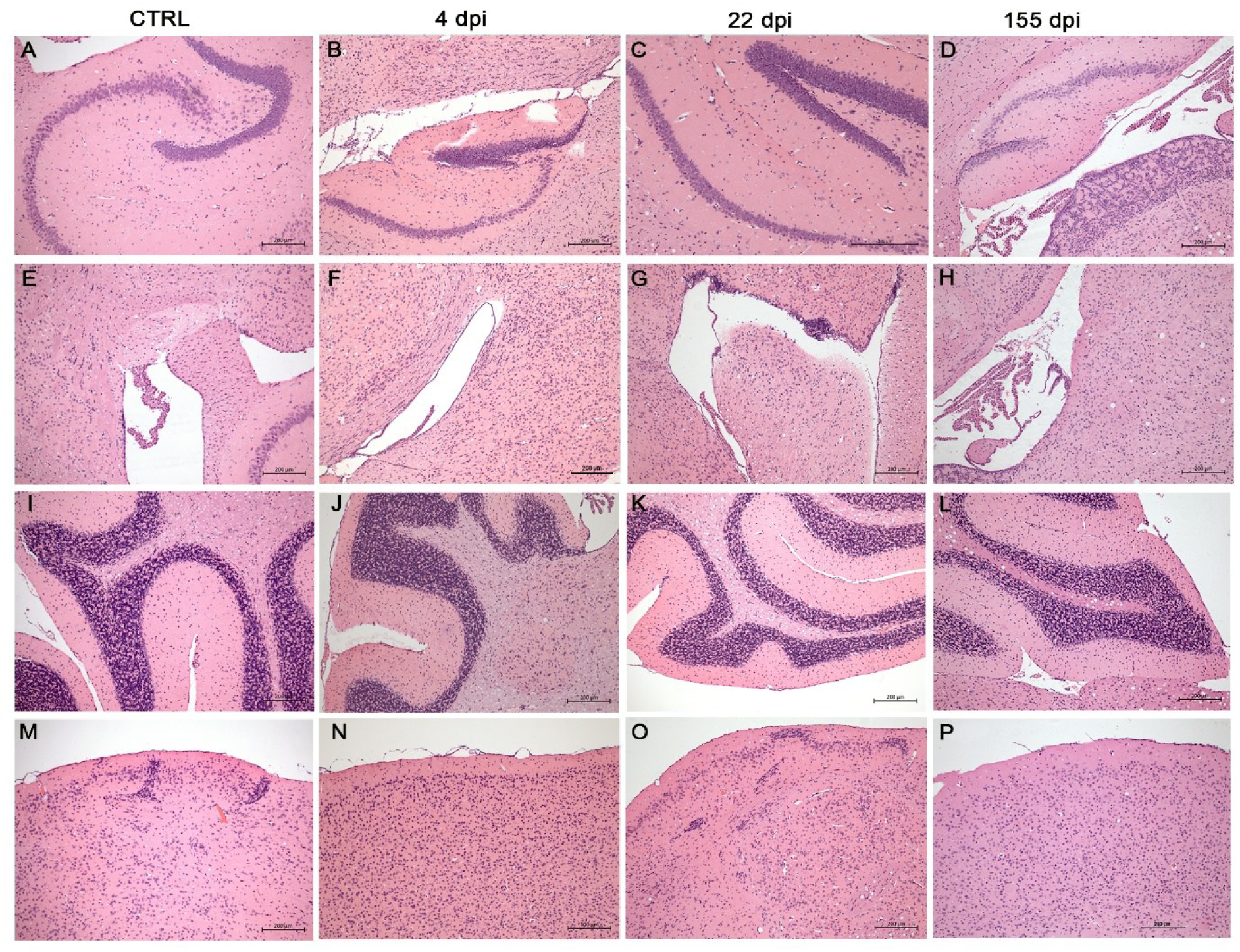
Effect of non-severe experimental plasmodial infection on brain morphology by hematoxylin-eosin staining. Histological sections of the hippocampus **(A-D)**, ventricular region with choroid plexus **(E-H)**, cerebellum **(I-L)** and cortex **(M-P)** with absence of inflammatory cells and typical architecture in control C57BL/6 mice (uninfected of cohort at D4) **(A, E, I, M)** and, 4 **(B, F, J, N)**, 22 **(C, G, K, O)** and 155 days post *Pb*A-infection **(D, H, L, P)**, stained with hematoxylin-eosin showed no morphological differences between groups (n=5). Scale bars = 200µm.

The Bielschowsky silver impregnation was performed to assess neuronal ramifications, and neuronal lesions (axonal injury, demyelination or neurodegeneration). Both control and infected mice showed no structural difference across all brain regions analyzed (**Figure 9**). Same as HE, the images were analyzed at higher magnifications, however, represented at low magnification to illustrate the brain regions.

**Figure 9.**
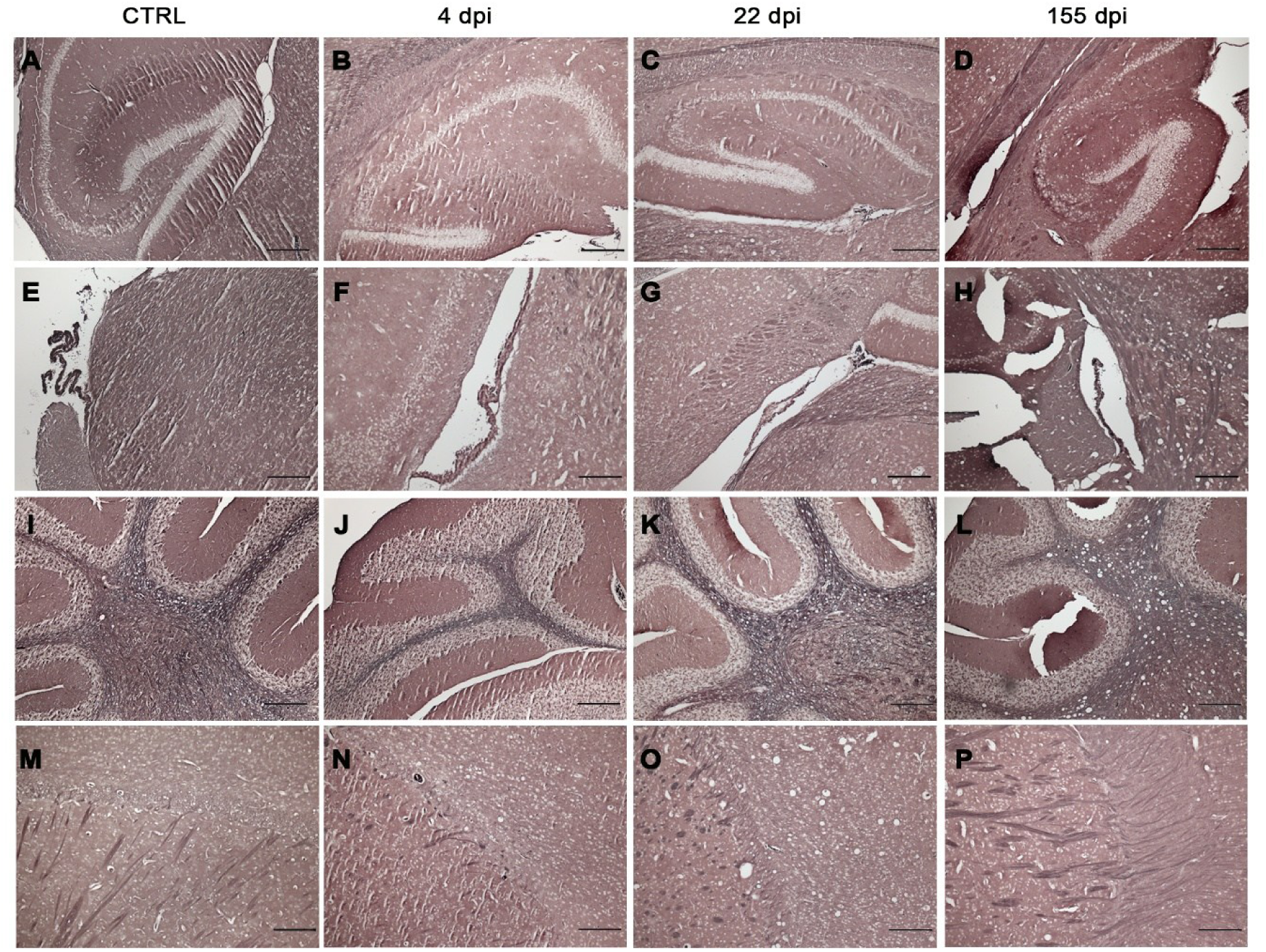
Effect of non-severe experimental plasmodial infection on brain morphology by Silver impregnation. Histological sections of the hippocampus **(A-D)**, ventricular region with choroid plexus **(E-H)**, cerebellum **(I-L)** and cortex **(M-P)**, without morphological changes in the different brain regions of control C57BL/6 mice (uninfected of cohort at D4) **(A, E, I, M)** and, 4 **(B, F, J, N)**, 22 **(C, G, K, O)** and 155 days post *Pb*A-infection **(D, H, L, P)**, using the Bielschowsky method (silver impregnation, showed no neuronal differences between groups) (n=3). Scale bars = 200 µm.

### Brain cellular and molecular alterations accompany sustained systemic inflammation in non-severe experimental malaria

To characterize the neuroinflammatory profile, hippocampal Th1, Th2, Th17 cytokine levels were measured by flow cytometry and ELISA at the maximum reached mean parasitemia (∼2.5%) in our 4 dpi protocol and after CQ treatment at 22 dpi. By flow cytometry, infected mice did not display different profiles from those of uninfected mice, at both 4 and 22 dpi (**Figure 10A)**. In the same way, no alterations were observed, by ELISA, in hippocampal TNF-α, IL-1β and IL-10 levels in infected mice (4 and 22 dpi) (**Figure 10B**). Hippocampal gene expression levels were analyzed by RT-qPCR at 4 dpi and an increased gene expression of hippocampal IL-1β levels was detected in infected mice (**Figure 10C)**.

**Figure 10.**
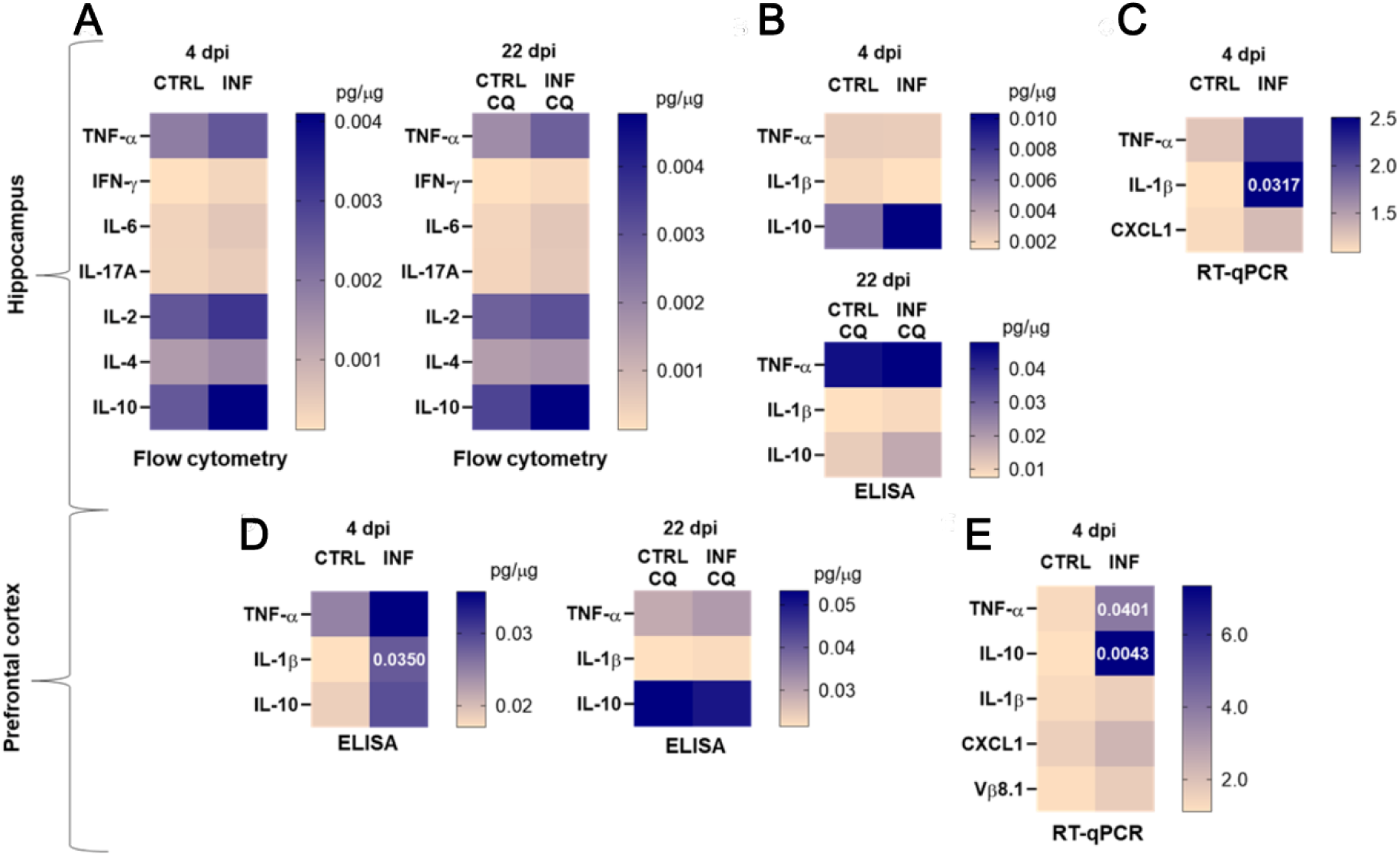
Effect of non-severe experimental plasmodial infection on brain molecular gene and protein expressions. Heatmaps represent a differentiated gene and protein expression profile in infected animals compared to the control. Hippocampal cytokine (TNF-α, IFN-ɣ, IL-6, IL-17A, IL-2, IL-4, IL-10 and IL-1β) levels by flow cytometry (n=4-9) **(A)** and ELISA (n=3-8) **(B)** of mice at 4 and 22 days post-*Plasmodium berghei* ANKA infection and gene expression (TNF-α, IL-1β and CXCL1) levels by RT-qPCR (n=2-5) **(C)** at 4 dpi were measured. Prefrontal cortex cytokine (TNF-α, IL-1β and IL-10) levels by ELISA (n=4-10) **(D)** at 4 and 22 dpi and gene expression (TNF-α, IL-10, IL-1β, CXCL1 and Vβ8.1) levels by RT-qPCR (n=5-9) **(E)** at 4 dpi are represented in heatmaps. CTRL: uninfected; INF: infected; CTRL-CQ: uninfected and treated with CQ; INF-CQ: infected and treated with CQ. Mann-Whitney test. p-values <0.05 in the heatmaps.

Increased levels of IL-1β were detected by ELISA in the prefrontal cortex of infected mice 4 dpi. After CQ-treatment, no difference was detected in cytokine levels at 22 dpi (**Figure 10D**). Although no significant differences were found in the gene expression of IL-1β, CXCL1 and Vβ8.1 in the prefrontal cortex at infected mice 4 dpi, an increase was recorded in the gene expression levels of TNF-α and IL-10 (**Figure 10E**).

Astrocytes are important cells in the metabolic maintenance of neurons and are functionally sensitive to alterations in contexts such as systemic inflammation (Guo *et al*., 2024). The hippocampal glial fibrillary acidic protein (GFAP) was analyzed in brain immunofluorescence slices (**Figure 11**). Infected mice showed a mild thickening of branched process in GFAP^+^ cells and the antimalarial CQ treatment reduced the GFAP density in the *cornu Ammonis* 1 (CA1) and dentate gyrus (DG) regions of the hippocampus (**Figure 11A, B**). These same profiles were observed when analyzing endpoints, average branch lengths, branch numbers and longest shortest path (**Figure 11C**). Mice uninfected and treated with CQ were analyzed and no difference in GFAP immunoreactivity and skeleton morphology in the hippocampus was detected when compared to material from uninfected and PBS-treated mice (**Figure S2**).

**Figure 11.**
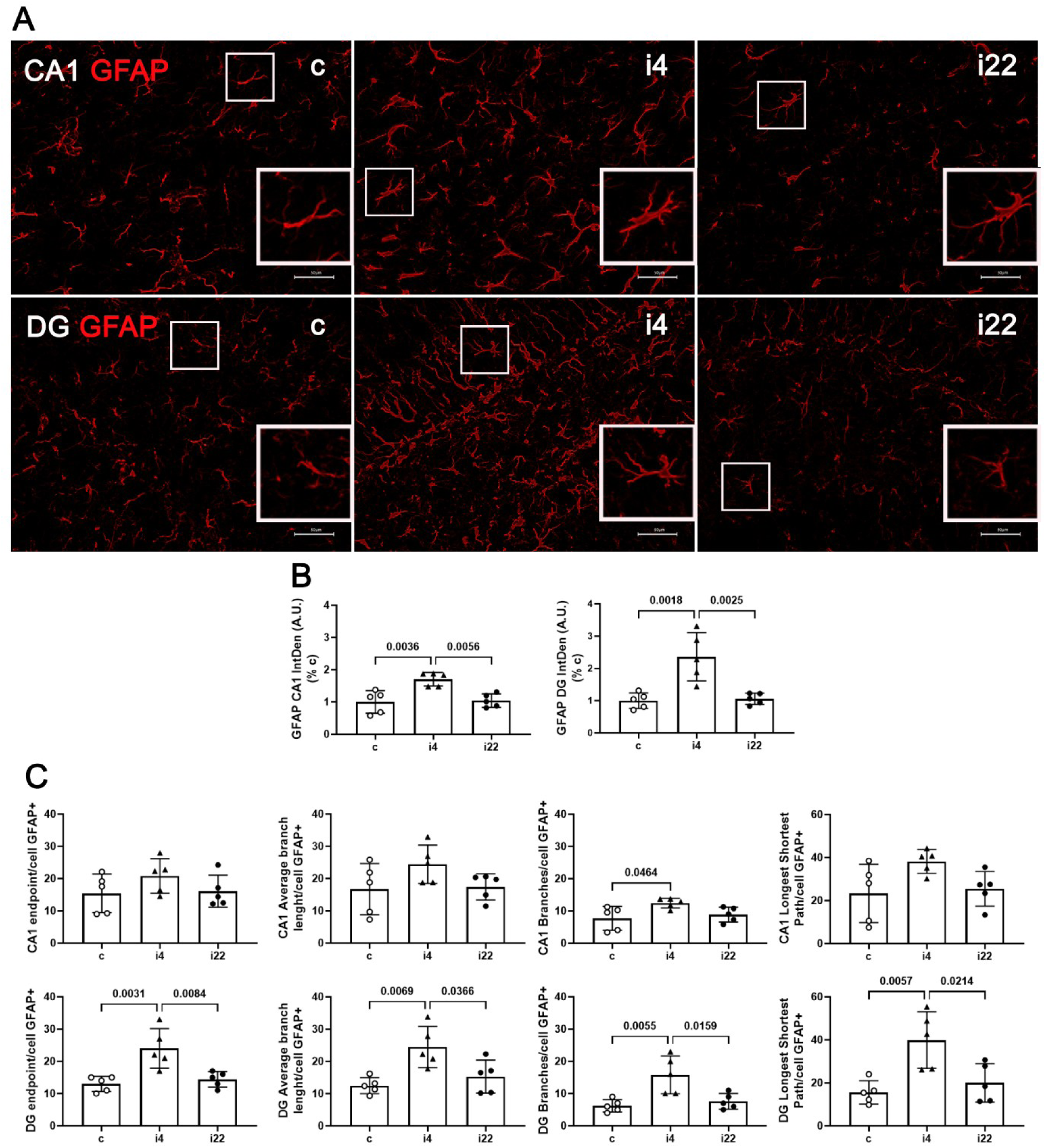
Effect of non-severe plasmodial infection on immunoreactivity of hippocampal astrocytes. Immunoreactivity of GFAP^+^ cells was detected in *cornu Ammonis* 1 (CA1) and dentate gyrus (DG) regions of the hippocampus of *Pb*A-infected and CQ-treated mice. Distinguishable photomicrograph (**A**) and quantification profile of integrated density (**B**) of GFAP immunoreactivity of CA1 and DG regions modulated by *Pb*A infection. **(C)** Cytoskeleton analysis of GFAP^+^ cells in hippocampal CA1 and DG regions. Columns represent mean ± standard deviation of uninfected control (c) treated with CQ, *Pb*A infected mice analyzed at 4dpi, before treatment (i4), as well *Pb*A-infected (i22) treated with CQ (n=5). Scale bars: 50 µm. The same scale bar at magnification is 25 µm. Data points are identified as individuals’ values. Columns represent mean ± S.D. One-way ANOVA/Bonferroni was used for comparison of c/i4, c/i22 and i4/i22. p-values <0.05 in the graphs.

Microglia are important cells in the maintenance of brain homeostasis and are also functionally sensitive to homeostasis alterations, such as infection (Chagas *et al*., 2020). The hippocampal ionized calcium-binding adaptor molecule 1 (Iba-1) protein was analyzed in brain immunofluorescence slices (**Figure 12**). No differences were detected in the immunoreactivity of Iba-1 cells in the experimental groups studied in both CA1 and DG regions of the hippocampus (**Figure 12A, B**). A reduction in endpoints, average branch length and branches per Iba-1^+^ cells were found in the CA1 region of the hippocampus of infected mice, suggesting a less complex branch phenotype of microglia, that persisted after CQ treatment in endpoints and branches (**Figure 12C**). No differences were detected in the skeleton analysis of DG region. Mice uninfected and treated with CQ showed no difference of Iba-1 immunoreactivity and skeleton morphology when compared to hippocampal brain slices of uninfected mice treated with vehicle PBS (**Figure S3**).

**Figure 12.**
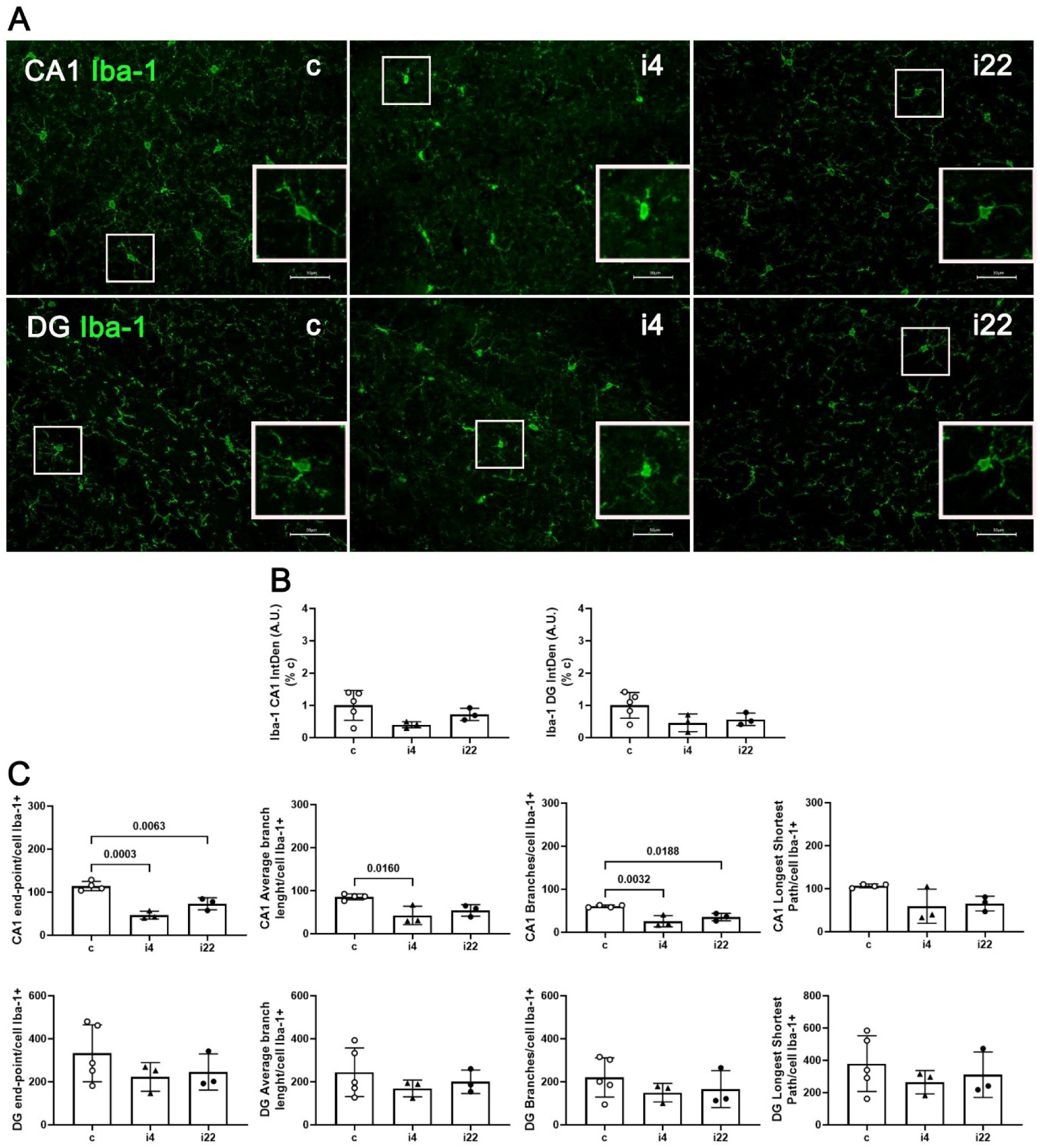
Effect of non-severe plasmodial infection on immunoreactivity of hippocampal Iba-1 cells. Immunoreactivity of Iba-1 cells was detected in the dentate gyrus (DG) and *cornu Ammonis* 1 (CA1) regions of the hippocampus of *Pb*A-infected and CQ-treated mice. Photomicrographs (**A**) and quantification profile of integrated density (**B**) of Iba-1 immunoreactivity of DG and CA1 region modulated by *Pb*A infection. (**C**) Cytoskeleton analysis of Iba-1^+^ cells in hippocampal DG and CA1 regions. Columns represent mean ± standard deviation of uninfected control (c) and *Pb*A-infected (i) mice treated with CQ, and *Pb*A-infected (i4) (n=3-5). Scale bars: 50 µm. The same scale bar at magnification is 25 µm. Data points are identified as individuals’ values. Columns represent mean ± S.D. One-way ANOVA/Bonferroni was used for comparison of c/i4, c/i22 and i4/i22. p-values <0.05 in the graphs.

## DISCUSSION

Approximately half of the world’s population is at risk for malaria, a potentially severe disease for around the 98% of cases on the planet that are caused by *P. falciparum* (WHO, 2024). In addition, malaria is considered an important cause of cognitive and behavioral alterations, impaired executive function and poor academic performance — in both severe and nSM, as well as in asymptomatic plasmodial infections (Vitor-Silva *et al*., 2009, Pessoa *et al*., 2022; Johnson *et al*., 2024), what correspond to an additional burden of the disease The pathophysiological events are poorly understood, and cellular and neurochemical studies are welcome (de Lima *et al*., 2025).

In this study, we conducted cellular, molecular, histological and behavioral analyses using an adapted nSEM model, and we established that non-severe *Pb*A infection displays a clear systemic proinflammatory profile accompanied by hippocampal and prefrontal cortex molecular inflammatory alterations. This conjunction of systemic and neuroinflammatory responses, including glial cell activation, may underlie the observed long-lasting cognitive impairment.

In this study, we have shown that *Pb*A infection did not affect the habituation memory, in accordance with our previous studies (de Sousa *et al*., 2018, 2021; Rosa-Gonçalves *et al*., 2022b). Recognition memory, a long-term episodic memory associated with memories of facts, events and people, is consolidated in the hippocampus that has, therefore, together with the cortex, an important role in its formation (Mendez *et al*., 2015). We found that it is impaired in *Pb*A-infected / treated mice, as previously shown by the poor performance in the ORT (de Sousa *et al*., 2018, 2021; Rosa-Gonçalves *et al*., 2022b). In this study we originally described that *Pb*A infection can also affect the learning acquisition process, as shown by the poor performance in the Object Location test, compromising the memory consolidation process (Vogel-Ciernia *et al*., 2014).

Environmental factors have been identified as risk factors for cognitive impairment (Da Ros *et al*., 2025). Infectious diseases such as sepsis, influenza, dengue, syphilis, toxoplasmosis, Chagas disease, COVID-19 and malaria have been associated with an increased risks of cognitive impairment that may contribute to the development and progression of neuropsychiatric and neurodegenerative disorders (da Silva Chagas *et al*., 2021; Barbosa-Silva *et al*., 2021). Neuroinflammation, which can also result from non-infectious inflammatory conditions, appears to be a common mechanism for developing these sequelae (Barbosa-Silva *et al*., 2021; Pascoal *et al*., 2021). The SARS-CoV-2 infection, for instance, is associated with persistent effects on attention and memory (Baig, 2022). Some pathogens share mechanisms that can lead to nervous system damage or neuronal degeneration, the pro-inflammatory cytokines impairing brain function and cognition (Kipnis *et al*., 2012; Schilling *et al*., 2021; Hernandez-Ruiz *et al*., 2022).

The dynamics of the cytokine levels observed in serum and spleen, in the present work, confirm that non-severe *Pb*A-infection can promote important systemic inflammatory changes that persist after antimalarial treatment and can contribute to the long-lasting cognitive and behavioral sequelae (Rosa-Gonçalves *et al*. 2022b). Johnson *et al*. (2024) identified cognitive deficit in school children mediated by subclinical inflammation in asymptomatic *P. falciparum* infection and TNF-α was the most consistent cytokine contributor to the outcome. High frequencies of total CD4^+^ and CD8^+^ T cells in the spleen were reported in *Pb*A-infected and treated mice and could be responsible for a large part of cytokine production in the inflammatory process of infected animals (de Sousa *et al*., 2021).

Dark spleen, characteristic of hemozoin pigmentation, was observed at 84 days after the end of CQ treatment (de Sousa *et al*., 2021) and, in our study, about five months after a successfully treated single episode of nSEM. The persistent pigmentation may start contributing to the activation of the immune system at the spleen, a secondary lymphoid organ where the initial immune response to infection occurs with pRBC being cleared from the circulation. Even after being taken up by neutrophils and macrophages, hemozoin can contribute to the increase of pathogen- and damage-associated molecular patterns (PAMPs and DAMPs) that are subsequently released into circulation and can stimulate monocytes and macrophages to produce neuroinflammatory cytokines such as IL-1ꞵ (Dostert *et al*., 2009; Griffith *et al*., 2009; Velagapudi *et al*., 2019; Pham *et al*., 2020; Chandana *et al*., 2020). Hemozoin in the early stage of infection is an important inflammatory factor in the spleen, but in the late stage, it may be suppressive due to its accumulation, paralyzing immune cells, what agrees with no evident cytokine response at 155 dpi (Urban *et al*., 2006).

We investigated the brain histopathology and none of the signs classically reported in CM, as demyelination, axonal injury, blood-brain barrier (BBB) disruption, inflammatory infiltrates, edema, hemorrhages, vascular congestion, deposits of amorphous material and necrosis were found in any brain region analyzed using hematoxylin-eosin or Bielschowsky silver staining (Milner *et al*., 2014; Strangward *et al*., 2017). The presence of moderate intravascular infiltrates and hemorrhagic areas has been reported at 5 dpi, the time when experimental CM begins, while no changes were found at 3 dpi (Lacerda-Queiroz *et al*., 2010; de Miranda *et al*., 2011). However, a subtle adherence of leukocytes and minimal edema were evidenced four days post-*Pb*A infection, although clinical signs of CM syndrome and BBB disruption were observed only at 6 dpi (Potter *et al*., 1999; Lima *et al*., 2020). As hypothesized by de Souza *et al*. (2018), physiological and molecular changes may occur in the absence of histopathological damage, resulting in cognitive impairment in experimental cerebral malaria (ECM). Mohanty *et al*. (2022) observed a subtle presence of edema in patients with severe non-cerebral malaria, using magnetic resonance imaging, due to a slight increase in systemic vascular permeability.

Our results suggest that the BBB function could be compromised at the initial stages of malaria, consistent with the detection of subtle changes in BBB permeability using Tracer-653 at the 4^th^ dpi in the same model (Freeman *et al*., 2016). The brain / peripheral immune system crosstalk displays through complex anatomical routes (Kovacs *et al*., 2025). Lymphoid organs like the spleen and lymph nodes are central hubs for immune cell activation. When these organs are inflamed — such as during a systemic infection like malaria — they can release proinflammatory cytokines (e.g., IL-1β, TNF-α, IL-6) into circulation. These circulating cytokines can affect the central nervous system in different ways (a) via the BBB, when its permeability is altered, even slightly, in response to infection or injury. The BBB can also have its endothelial cells activated to produce secondary signals, promoting the access of immune molecules, pathogens, and neurotoxic components to the brain, in addition to stimulating microglia and astrocytes, and other glial cells, which may amplify central pro-inflammatory signaling. The glial response leads to a positive feedback loop, exacerbating neuroinflammation (Daneman and Prat, 2015); (b) Another way to systemic inflammation accesses the CNS is by signaling through circumventricular organs that do not have a BBB (Kovacs *et al*., 2025; Daneman and Prat, 2015).

We detected an increase of gene and protein expressions of pro-inflammatory cytokines, such as IL-1ꞵ and TNF-ɑ, in cognitive relevant brain regions of infected mice, at the 4^th^ dpi, corroborating the findings of de Miranda *et al*. (2011), who showed high brain levels of IL-1β and TNF-α at 5 dpi, associated with anxiety-like behavior in ECM (Miranda *et al*., 2011). The increased gene expression of TNF-α and IL-10 in prefrontal cortex probably reflects an autoregulation of cytokines. Elevated levels of TNF-ɑ in cerebrospinal fluid of children with CM were also associated with long-term cognitive sequelae (Shabani *et al*., 2017). In experimental CM, IL-1ꞵ released by microglia, mediated by the IL-33/ST2 pathway, impairs neurogenesis and cognition (Reverchon *et al*., 2017). The upregulation of chemokines from brain microvascular endothelium and astrocytes, which potentiate the entry of neurotoxic neutrophils and neuronal loss, was found to contribute to cognitive impairments in Lyme disease (Brissette *et al*., 2013), and similar events could operate in malaria cognitive sequelae.

The hippocampal DG and CA1 regions play a critical role in memory formation and are often compromised in diseases that affect cognition, such as Alzheimer’s and Parkinson’s diseases (Sahin *et al*., 2024; Fixemer *et al*., 2025). These regions have been identified as having an important association for spatial memory such as the revealed in the object location test in mice (Assini *et al*., 2009; Frechou *et al*., 2024). It has been suggested that the DG, particularly its new neurons generated via adult neurogenesis, is involved in memory acquisition and recall (Aasebø *et al*., 2018). In our study, we analyzed immunoreactivity for GFAP, an astrocyte marker found gradually elevated in the ECM (Oelschlegel *et al*., 2024), in the DG and CA1 of the hippocampus and observed characteristics suggestive of astrocyte reactivity in response to infection, with the effect particularly marked in DG. Notably, the DG of the hippocampus — a region critically involved in memory acquisition and the encoding of novel information — has been shown to be particularly vulnerable to systemic inflammatory insults. According to Habbas *et al*. (2015), inflammation-induced changes in astrocyte activity within the DG are associated with impaired performance in hippocampal-dependent memory tasks, underscoring the DG’s essential role in early memory processing. The cognitive damage detected in our experimental model could, therefore, be related to astrocytic dysfunction following early BBB permeability alterations, particularly affecting regions such as the DG that are central to memory formation.

Altered microglial states have also been identified in brain diseases when microglia respond to microenvironment challenges (Chagas *et al*., 2020; Paolicelli *et al*., 2022). In our study, we evidenced a subtle microglial profile, as revealed by significant reduction per Iba1^+^ cell in endpoints, average branch length and branches in CA1 region of infected mice. Although Iba1^+^ does not directly distinguish microglial phenotypes, the expression of this molecule reflects morphological and cell density changes, which are associated with microglial functional phenotypes. Thus, changes in Iba1^+^ expression may provide indirect indicators of microglial functional changes. In this context, stable modifications in histones and DNA methylation occur, altering microglial gene expression in the long-term (Lima *et al*., 2022). The alteration in microglial phenotype, such as the branch retraction and loss of complexity we detected in the early stages of plasmodial infection, were still detected after antimalarial treatment and might be suggestive of persistent inflammatory response. Our data agree with those of Medana *et al*. (2002), identifying microglia morphological alterations in the early stages of plasmodial infection, before BBB disruption and brain pathology. Thus, microglia, that may appear morphologically homeostatic, could be primed by exposure to plasmodial infection and be molecularly and functionally altered (Lima *et al*., 2022). It became evident from our results that, even treated at its early stages, *Pb*A infection can promote a long-lasting shift from the homeostatic microglial phenotype and it is, therefore, possible that the functional and morphological changes in these cells is involved in the genesis of the long-term cognitive deficits observed in our study.

The specific functions of glial cells need to be addressed in the context of their multiple functionality states in future studies in nSEM, and since the present study was conducted on females, studies investigating changes in males are still required before any definitive conclusion is reached.

In conclusion, nSM caused by *Pb*A can promote systemic inflammation, cellular and molecular alterations in the brain, especially in regions of cognitive importance (hippocampus and prefrontal cortex), with involvement of glial cells and mild neuroinflammation. Such events can be associated with cognitive sequelae detected at different times, even in the absence of detectable morphological changes in hematoxylin-eosin and Bielschowsky staining methods.

## CRediT AUTHORSHIP CONTRIBUTION STATEMENT

**PRG & BNSS:** Writing – Original Draft Preparation, Writing – Review & Editing, Visualization, Validation, Methodology, Investigation, Formal Analysis, Conceptualization; **MNL:** Writing – Review & Editing, Formal Analysis, Investigation, Visualization; **LSC:** Writing – Review & Editing, Investigation, Visualization; **LPS:** Writing – Review & Editing, Investigation; **LFGC:** Writing – Review & Editing, Methodology, Investigation; **IJS:** Writing – Review & Editing, Investigation; **JPRS**: Writing – Review & Editing, Investigation; **CCAE:** Writing – Review & Editing, Investigation; **MSN:** Writing – Review & Editing, Investigation; **GLW**: Writing – Review & Editing, Formal Analysis; **FLRG:** Writing – Review & Editing, Investigation, Supervision; **PRRT:** Writing – Review & Editing, Investigation, Visualization, Supervision; **LRPT:** Writing – Review & Editing, Investigation, Supervision; **TMG:** Writing – Review & Editing, Methodology, Resources, Supervision; **RFA:** Writing – Review & Editing, Methodology, Conceptualization, Supervision; **MPM, CAS & CTDR:** Writing – Original Draft Preparation, Writing – Review & Editing, Investigation, Conceptualization, Funding Acquisition, Methodology, Project Administration, Resources, Supervision.

## ACKNOWLEDGMENTS

The authors thank the Animal Facilities team of *Centro de Experimentação Animal* (*IOC, Fiocruz*) for help with animal care and experiments and thank the *Instituto de Ciência e Tecnologia em Biomodelos* (*Fiocruz*) for providing the animals. We are grateful to Dr. Rudimar Frozza for teaching PRG to dissect the hippocampus and prefrontal cortex and to Dr. Otacilio Moreira for suggesting data representation in heat maps. We thank the Multi-user Research Facility of Flow Cytometry, *Instituto Oswaldo Cruz*, *Fiocruz*, Rio de Janeiro, Brazil. We also thank Cynthia Cascabulho and Bárbara du Rocher for help with flow cytometry analyses, Poliana Capucho for helping with brain’s processing protocol, and teaching and helping with the cryostat with Samuel Horita. We also thank Farmanguinhos (*Fiocruz*) through Juliana Johanson for the donation of the antimalarial CQ. We are grateful to *Laboratório de Pesquisas sobre o Timo* for sharing reagents and equipment.

## REFERENCES

Aasebø IEJ, Kasture AS, Passeggeri M, Tashiro A. A behavioral task with more opportunities for memory acquisition promotes the survival of new neurons in the adult dentate gyrus. Sci Rep. 2018 May 9;8(1):7369. doi: 10.1038/s41598-018-25331-w.

Allen Institute for Brain Science (2004). Allen Mouse Brain Atlas [dataset]. Available from mouse.brain-map.org. Allen Institute for Brain Science (2011). Allen Reference Atlas – Mouse Brain [brain atlas]. Available from atlas.brain-map.org.

Assini FL, Duzzioni M, Takahashi RN. Object location memory in mice: pharmacological validation and further evidence of hippocampal CA1 participation. Behav Brain Res. 2009 Dec 1;204(1):206–11. doi: 10.1016/j.bbr.2009.06.005.

Baig AM. Counting the neurological cost of COVID-19. Nat Rev Neurol. 2022 Jan;18(1):5–6. doi: 10.1038/s41582-021-00593-7.

Barbosa-Silva MC, Lima MN, Battaglini D, Robba C, Pelosi P, Rocco PRM, Maron-Gutierrez T. Infectious disease-associated encephalopathies. Crit Care. 2021 Jul 6;25(1):236. doi: 10.1186/s13054-021-03659-6.

Brissette CA, Kees ED, Burke MM, Gaultney RA, Floden AM, Watt JA. The multifaceted responses of primary human astrocytes and brain microvascular endothelial cells to the Lyme disease spirochete, Borrelia burgdorferi. ASN Neuro. 2013 Aug 16;5(3):221–9. doi: 10.1042/AN20130010.

Caputo LFG, Gitirana LB, Manso PPA. 2011. Técnicas histológicas, in: Molinaro EM, Caputo LFG, Amendoeira RMR (Orgs.), Conceitos e Métodos para a Formação de Profissionais em Laboratórios de Saúde. Vol 2. Editora Fiocruz, Rio de Janeiro, pp. 146–149.

Chagas LDS, Sandre PC, Ribeiro E Ribeiro NCA, Marcondes H, Oliveira Silva P, Savino W, Serfaty CA. Environmental Signals on Microglial Function during Brain Development, Neuroplasticity, and Disease. Int J Mol Sci. 2020 Mar 19;21(6):2111. doi: 10.3390/ijms21062111.

Chagas LDS, Trindade P, Gomes ALT, Mendonça HR, Campello-Costa P, Faria Melibeu ADC, Linden R, Serfaty CA. Rapid plasticity of intact axons following a lesion to the visual pathways during early brain development is triggered by microglial activation. Exp Neurol. 2019 Jan;311:148–161. doi: 10.1016/j.expneurol.2018.10.002.

Chandana M, Anand A, Ghosh S, Das R, Beura S, Jena S, Suryawanshi AR, Padmanaban G, Nagaraj VA. Malaria parasite heme biosynthesis promotes and griseofulvin protects against cerebral malaria in mice. Nat Commun. 2022 Jul 12;13(1):4028. doi: 10.1038/s41467-022-31431-z. Erratum in: Nat Commun. 2024 Jun 25;15(1):5363. doi: 10.1038/s41467-024-49627-w.

Coban C, Lee MSJ, Ishii KJ. Tissue-specific immunopathology during malaria infection. Nat Rev Immunol. 2018 Apr;18(4):266–278. doi: 10.1038/nri.2017.138.

Conroy AL, Datta D, John CC. What causes severe malaria and its complications in children? Lessons learned over the past 15 years. BMC Med. 2019 Mar 7;17(1):52. doi: 10.1186/s12916-019-1291-z.

Da Ros LU, Borelli WV, Aguzzoli CS, De Bastiani MA, Schilling LP, Santamaria-Garcia H, Pascoal TA, Rosa-Neto P, Souza DO, da Costa JC, Ibañez A, Suemoto CK, Zimmer ER. Social and health disparities associated with healthy brain ageing in Brazil and in other Latin American countries. Lancet Glob Health. 2025 Feb;13(2):e277–e284. doi: 10.1016/S2214-109X(24)00451-0.

da Silva Chagas L, Sandre PC, de Velasco PC, Marcondes H, Ribeiro E Ribeiro NCA, Barreto AL, Alves Mauro LB, Ferreira JH, Serfaty CA. Neuroinflammation and Brain Development: Possible Risk Factors in COVID-19-Infected Children. Neuroimmunomodulation. 2021;28(1):22–28. doi: 10.1159/000512815.

Daneman R, Prat A. The blood-brain barrier. Cold Spring Harb Perspect Biol. 2015 Jan 5;7(1):a020412. doi: 10.1101/cshperspect.a020412.

de Lima RMS, Leão LKR, Martins LC, Passos ADCF, Batista EJO, Herculano AM, Oliveira KRHM. Unveiling new perspectives about the onset of neurological and cognitive deficits in cerebral malaria: exploring cellular and neurochemical mechanisms. Front Cell Infect Microbiol. 2025 Feb 6;15:1506282. doi: 10.3389/fcimb.2025.1506282.

de Miranda AS, Brant F, Vieira LB, Rocha NP, Vieira ÉLM, Rezende GHS, de Oliveira Pimentel PM, Moraes MFD, Ribeiro FM, Ransohoff RM, Teixeira MM, Machado FS, Rachid MA, Teixeira AL. A Neuroprotective Effect of the Glutamate Receptor Antagonist MK801 on Long-Term Cognitive and Behavioral Outcomes Secondary to Experimental Cerebral Malaria. Mol Neurobiol. 2017 Nov;54(9):7063–7082. doi: 10.1007/s12035-016-0226-3.

de Miranda AS, Lacerda-Queiroz N, de Carvalho Vilela M, Rodrigues DH, Rachid MA, Quevedo J, Teixeira AL. Anxiety-like behavior and proinflammatory cytokine levels in the brain of C57BL/6 mice infected with *Plasmodium berghei* (strain ANKA). Neurosci Lett. 2011 Mar 24;491(3):202–6. doi: 10.1016/j.neulet.2011.01.038.

de Sousa LP, de Almeida RF, Ribeiro-Gomes FL et al. Long-term effect of uncomplicated *Plasmodium berghei* ANKA malaria on memory and anxiety-like behaviour in C57BL/6 mice. Parasit Vectors. 2018 Mar 20;11(1):191. doi: 10.1186/s13071-018-2778-8.

de Sousa LP, Ribeiro-Gomes FL, de Almeida RF et al. Immune system challenge improves recognition memory and reverses malaria-induced cognitive impairment in mice. Sci Rep. 2021 Jul 21;11(1):14857. doi: 10.1038/s41598-021-94167-8.

Dostert C, Guarda G, Romero JF, Menu P, Gross O, Tardivel A, Suva ML, Stehle JC, Kopf M, Stamenkovic I, Corradin G, Tschopp J. Malarial hemozoin is a Nalp3 inflammasome activating danger signal. PLoS One. 2009 Aug 4;4(8):e6510. doi: 10.1371/journal.pone.0006510.

Fernando SD, Rodrigo C, Rajapakse S. The ‘hidden’ burden of malaria: cognitive impairment following infection. Malar J. 2010 Dec 20;9:366. doi: 10.1186/1475-2875-9-366.

Fixemer S, Miranda de la Maza M, Hammer GP, Jeannelle F, Schreiner S, Gérardy JJ, Boluda S, Mirault D, Mechawar N, Mittelbronn M, Bouvier DS. Microglia aggregates define distinct immune and neurodegenerative niches in Alzheimer’s disease hippocampus. Acta Neuropathol. 2025 Feb 15;149(1):19. doi: 10.1007/s00401-025-02857-8.

Frechou MA, Martin SS, McDermott KD, Huaman EA, Gökhan Ş, Tomé WA, Coen-Cagli R, Gonçalves JT. Adult neurogenesis improves spatial information encoding in the mouse hippocampus. Nat Commun. 2024 Jul 30;15(1):6410. doi: 10.1038/s41467-024-50699-x.

Freeman BD, Martins YC, Akide-Ndunge OB, Bruno FP, Wang H, Tanowitz HB, Spray DC, Desruisseaux MS. Endothelin-1 Mediates Brain Microvascular Dysfunction Leading to Long-Term Cognitive Impairment in a Model of Experimental Cerebral Malaria. PLoS Pathog. 2016 Mar 31;12(3):e1005477. doi: 10.1371/journal.ppat.1005477.

Gazzinelli RT, Kalantari P, Fitzgerald KA, Golenbock DT. Innate sensing of malaria parasites. Nat Rev Immunol. 2014 Nov;14(11):744–57. doi: 10.1038/nri3742.

Griffith JW, Sun T, McIntosh MT, Bucala R. Pure Hemozoin is inflammatory in vivo and activates the NALP3 inflammasome via release of uric acid. J Immunol. 2009 Oct 15;183(8):5208–20. doi: 10.4049/jimmunol.0713552.

Guo Q, Gobbo D, Zhao N, Zhang H, Awuku NO, Liu Q, Fang LP, Gampfer TM, Meyer MR, Zhao R, Bai X, Bian S, Scheller A, Kirchhoff F, Huang W. Adenosine triggers early astrocyte reactivity that provokes microglial responses and drives the pathogenesis of sepsis-associated encephalopathy in mice. Nat Commun. 2024 Jul 27;15(1):6340. doi: 10.1038/s41467-024-50466-y. Erratum in: Nat Commun. 2024 Sep 18;15(1):8200. doi: 10.1038/s41467-024-52497-x.

Habbas S, Santello M, Becker D, Stubbe H, Zappia G, Liaudet N, Klaus FR, Kollias G, Fontana A, Pryce CR, Suter T, Volterra A. Neuroinflammatory TNFα Impairs Memory via Astrocyte Signaling. Cell. 2015 Dec 17;163(7):1730–41. doi: 10.1016/j.cell.2015.11.023.

Hernandez-Ruiz V, Letenneur L, Fülöp T, Helmer C, Roubaud-Baudron C, Avila-Funes JA, Amieva H. Infectious diseases and cognition: do we have to worry? Neurol Sci. 2022 Nov;43(11):6215–6224. doi: 10.1007/s10072-022-06280-9.

Izquierdo I, Medina JH. Memory formation: the sequence of biochemical events in the hippocampus and its connection to activity in other brain structures. Neurobiol Learn Mem. 1997 Nov;68(3):285–316. doi: 10.1006/nlme.1997.3799. PMID: 9398590.

Johnson AE, Upadhye A, Knight V, Gaskin EL, Turnbull LB, Ayuku D, Nyalumbe M, Abuonji E, John CC, McHenry MS, Tran TM, Ayodo G. Subclinical Inflammation in Asymptomatic Schoolchildren With Plasmodium falciparum Parasitemia Correlates With Impaired Cognition. J Pediatric Infect Dis Soc. 2024 May 30;13(5):288–296. doi: 10.1093/jpids/piae025.

Kipnis J, Gadani S, Derecki NC. Pro-cognitive properties of T cells. Nat Rev Immunol. 2012 Sep;12(9):663–9. doi: 10.1038/nri3280.

Kovacs M, Dominguez-Belloso A, Ali-Moussa S, Deczkowska A. Immune control of brain physiology. Nat Rev Immunol. 2025 Jan 31. doi: 10.1038/s41577-025-01129-6.

Lacerda-Queiroz N, Rodrigues DH, Vilela MC, Miranda AS, Amaral DC, Camargos ER, Carvalho LJ, Howe CL, Teixeira MM, Teixeira AL. Inflammatory changes in the central nervous system are associated with behavioral impairment in Plasmodium berghei (strain ANKA)-infected mice. Exp Parasitol. 2010 Jul;125(3):271–8. doi: 10.1016/j.exppara.2010.02.002.

Lewis SM, Williams A, Eisenbarth SC. Structure and function of the immune system in the spleen. Sci Immunol. 2019 Mar 1;4(33):eaau6085. doi: 10.1126/sciimmunol.aau6085.

Lima MN, Oliveira HA, Fagundes PM et al. Mesenchymal stromal cells protect against vascular damage and depression-like behavior in mice surviving cerebral malaria. Stem Cell Res Ther. 2020 Aug 26;11(1):367. doi: 10.1186/s13287-020-01874-6.

Lima MN, Barbosa-Silva MC, Maron-Gutierrez T. Microglial Priming in Infections and Its Risk to Neurodegenerative Diseases. Front Cell Neurosci. 2022 Jun 15;16:878987. doi: 10.3389/fncel.2022.878987.

Medana IM, Day NP, Hien TT, Mai NT, Bethell D, Phu NH, Farrar J, Esiri MM, White NJ, Turner GD. Axonal injury in cerebral malaria. Am J Pathol. 2002 Feb;160(2):655–66. doi: 10.1016/S0002-9440(10)64885-7.

Mendez M, Arias N, Uceda S, Arias JL. c-Fos expression correlates with performance on novel object and novel place recognition tests. Brain Res Bull. 2015 Aug;117:16–23. doi: 10.1016/j.brainresbull.2015.07.004.

Milner DA Jr, Whitten RO, Kamiza S, Carr R, Liomba G, Dzamalala C, Seydel KB, Molyneux ME, Taylor TE. The systemic pathology of cerebral malaria in African children. Front Cell Infect Microbiol. 2014 Aug 21;4:104. doi: 10.3389/fcimb.2014.00104.

Mohanty S, Sahu PK, Pattnaik R et al. Evidence of Brain Alterations in Noncerebral Falciparum Malaria. Clin Infect Dis. 2022 Aug 24;75(1):11–18. doi: 10.1093/cid/ciab907.

Oelschlegel AM, Bhattacharjee R, Wenk P, Harit K, Rothkötter HJ, Koch SP, Boehm-Sturm P, Matuschewski K, Budinger E, Schlüter D, Goldschmidt J, Nishanth G. Beyond the microcirculation: sequestration of infected red blood cells and reduced flow in large draining veins in experimental cerebral malaria. Nat Commun. 2024 Mar 16;15(1):2396. doi: 10.1038/s41467-024-46617-w.

Paolicelli RC, Sierra A, Stevens B, Tremblay ME, Aguzzi A, Ajami B, Amit I, Audinat E, Bechmann I, Bennett M, Bennett F, Bessis A, Biber K, Bilbo S, Blurton-Jones M, Boddeke E, Brites D, Brône B, Brown GC, Butovsky O, Carson MJ, Castellano B, Colonna M, Cowley SA, Cunningham C, Davalos D, De Jager PL, de Strooper B, Denes A, Eggen BJL, Eyo U, Galea E, Garel S, Ginhoux F, Glass CK, Gokce O, Gomez-Nicola D, González B, Gordon S, Graeber MB, Greenhalgh AD, Gressens P, Greter M, Gutmann DH, Haass C, Heneka MT, Heppner FL, Hong S, Hume DA, Jung S, Kettenmann H, Kipnis J, Koyama R, Lemke G, Lynch M, Majewska A, Malcangio M, Malm T, Mancuso R, Masuda T, Matteoli M, McColl BW, Miron VE, Molofsky AV, Monje M, Mracsko E, Nadjar A, Neher JJ, Neniskyte U, Neumann H, Noda M, Peng B, Peri F, Perry VH, Popovich PG, Pridans C, Priller J, Prinz M, Ragozzino D, Ransohoff RM, Salter MW, Schaefer A, Schafer DP, Schwartz M, Simons M, Smith CJ, Streit WJ, Tay TL, Tsai LH, Verkhratsky A, von Bernhardi R, Wake H, Wittamer V, Wolf SA, Wu LJ, Wyss-Coray T. Microglia states and nomenclature: A field at its crossroads. Neuron. 2022 Nov 2;110(21):3458–3483. doi: 10.1016/j.neuron.2022.10.020.

Pascoal TA, Benedet AL, Ashton NJ, Kang MS, Therriault J, Chamoun M, Savard M, Lussier FZ, Tissot C, Karikari TK, Ottoy J, Mathotaarachchi S, Stevenson J, Massarweh G, Schöll M, de Leon MJ, Soucy JP, Edison P, Blennow K, Zetterberg H, Gauthier S, Rosa-Neto P. Publisher Correction: Microglial activation and tau propagate jointly across Braak stages. Nat Med. 2021 Nov;27(11):2048–2049. doi: 10.1038/s41591-021-01568-3.

Pessoa RC, Oliveira-Pessoa GF, Souza BKA, Sampaio VS, Pinto ALCB, Barboza LL, Mouta GS, Silva EL, Melo GC, Monteiro WM, Silva-Filho JH, Lacerda MVG, Baía-da-Silva DC. Impact of Plasmodium vivax malaria on executive and cognitive functions in elderlies in the Brazilian Amazon. Sci Rep. 2022 Jun 20;12(1):10361. doi: 10.1038/s41598-022-14175-0. Erratum in: Sci Rep. 2024 Jan 11;14(1):1120. doi: 10.1038/s41598-023-48865-0.

Pham TT, Lamb TJ, Deroost K, Opdenakker G, Van den Steen PE. Hemozoin in Malarial Complications: More Questions Than Answers. Trends Parasitol. 2021 Mar;37(3):226–239. doi: 10.1016/j.pt.2020.09.016.

Potter S, Chaudhri G, Hansen A, Hunt NH. Fas and perforin contribute to the pathogenesis of murine cerebral malaria. Redox Rep. 1999;4(6):333–5. doi: 10.1179/135100099101535070.

Prophet EB, Mills B, Arrington JB, Sobin LH. 1994. Armed Forces Institute of Pathology: Laboratory Methods in Histotechnology. American Registry of Pathology, Washington, DC.

Reverchon F, Mortaud S, Sivoyon M, Maillet I, Laugeray A, Palomo J, Montécot C, Herzine A, Meme S, Meme W, Erard F, Ryffel B, Menuet A, Quesniaux VFJ. IL-33 receptor ST2 regulates the cognitive impairments associated with experimental cerebral malaria. PLoS Pathog. 2017 Apr 27;13(4):e1006322. doi: 10.1371/journal.ppat.1006322.

Rosa-Gonçalves P, Ribeiro-Gomes FL, Daniel-Ribeiro CT. Malaria Related Neurocognitive Deficits and Behavioral Alterations. Front Cell Infect Microbiol. 2022a Feb 22;12:829413. doi: 10.3389/fcimb.2022.829413.

Rosa-Gonçalves P, de Sousa LP, Maia AB et al. Dynamics and immunomodulation of cognitive deficits and behavioral changes in non-severe experimental malaria. Front Immunol. 2022b Nov 24;13:1021211. doi: 10.3389/fimmu.2022.1021211.

Sahin S, Velioglu HA, Yulug B, Bayraktaroglu Z, Yildirim S, Hanoglu L. Parietal memory network and memory encoding versus retrieval impairments in PD-MCI patients: A hippocampal volume and cortical thickness study. CNS Neurosci Ther. 2024 Oct;30(10):e70062. doi: 10.1111/cns.70062.

Schilling S, Chausse B, Dikmen HO, Almouhanna F, Hollnagel JO, Lewen A, Kann O. TLR2- and TLR3-activated microglia induce different levels of neuronal network dysfunction in a context-dependent manner. Brain Behav Immun. 2021 Aug;96:80–91. doi: 10.1016/j.bbi.2021.05.013.

Schwarzer E, Turrini F, Ulliers D, Giribaldi G, Ginsburg H, Arese P. Impairment of macrophage functions after ingestion of Plasmodium falciparum-infected erythrocytes or isolated malarial pigment. J Exp Med. 1992 Oct 1;176(4):1033–41. doi: 10.1084/jem.176.4.1033. Erratum in: J Exp Med 1993 Mar 1;177(3):following 873.

Shabani E, Ouma BJ, Idro R, Bangirana P, Opoka RO, Park GS, Conroy AL, John CC. Elevated cerebrospinal fluid tumour necrosis factor is associated with acute and long-term neurocognitive impairment in cerebral malaria. Parasite Immunol. 2017 Jul;39(7):10.1111/pim.12438. doi: 10.1111/pim.12438.

Souza TL, Grauncke ACB, Ribeiro LR et al. Cerebral Malaria Causes Enduring Behavioral and Molecular Changes in Mice Brain Without Causing Gross Histopathological Damage. Neuroscience. 2018 Jan 15;369:66–75. doi: 10.1016/j.neuroscience.2017.10.043.

Ssemata AS, Nakitende AJ, Kizito S, Thomas MR, Islam S, Bangirana P, Nakasujja N, Yang Z, Yu Y, Tran TM, John CC, McHenry MS. Association of severe malaria with cognitive and behavioural outcomes in low- and middle-income countries: a meta-analysis and systematic review. Malar J. 2023 Aug 3;22(1):227. doi: 10.1186/s12936-023-04653-9.

Strangward P, Haley MJ, Shaw TN et al. A quantitative brain map of experimental cerebral malaria pathology. PLoS Pathog. 2017 Mar 8;13(3):e1006267. doi: 10.1371/journal.ppat.1006267.

Urban BC, Todryk S. Malaria pigment paralyzes dendritic cells. J Biol. 2006;5(2):4. doi: 10.1186/jbiol37.

Varo R, Crowley VM, Sitoe A, Madrid L, Serghides L, Kain KC, Bassat Q. Adjunctive therapy for severe malaria: a review and critical appraisal. Malar J. 2018 Jan 24;17(1):47. doi: 10.1186/s12936-018-2195-7.

Vitor-Silva S, Reyes-Lecca RC, Pinheiro TR, Lacerda MV. Malaria is associated with poor school performance in an endemic area of the Brazilian Amazon. Malar J. 2009 Oct 16;8:230. doi: 10.1186/1475-2875-8-230.

Vogel-Ciernia A, Wood MA. Examining object location and object recognition memory in mice. Curr Protoc Neurosci. 2014 Oct 8;69:8.31.1–17. doi: 10.1002/0471142301.ns0831s69.

Velagapudi R, Kosoko AM, Olajide OA. Induction of Neuroinflammation and Neurotoxicity by Synthetic Hemozoin. Cell Mol Neurobiol. 2019 Nov;39(8):1187–1200. doi: 10.1007/s10571-019-00713-4.

Watermeyer JM, Hale VL, Hackett F, Clare DK, Cutts EE, Vakonakis I, Fleck RA, Blackman MJ, Saibil HR. A spiral scaffold underlies cytoadherent knobs in Plasmodium falciparum-infected erythrocytes. Blood. 2016 Jan 21;127(3):343–51. doi: 10.1182/blood-2015-10-674002.

WHO - World Health Organization. World Malaria Report 2024. Geneva: World Health Organization; 2024.

Young K, Morrison H. Quantifying Microglia Morphology from Photomicrographs of Immunohistochemistry Prepared Tissue Using ImageJ. J Vis Exp. 2018 Jun 5;(136):57648. doi: 10.3791/57648.

