## Supplementary Material for "SUSTAINED SYSTEMIC AND NEUROINFLAMMATION IN COGNITIVE DYSFUNCTIONAL FEMALE MICE AFTER NON-SEVERE EXPERIMENTAL MALARIA"

**1 Supplementary Figures**

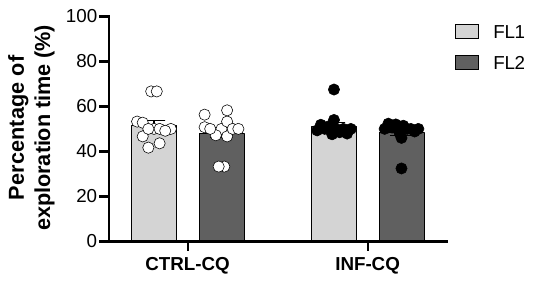

**Figure S1. Effect of non-severe plasmodial infection on recognition memory of mice in training session of object location test (OLT).** Time in objects at familiar locations in percentage (%) of uninfected and infected, both treated mice in training session, lasting 10 minutes in OLT cohort evaluated at 22 dpi. Values were manually timed with digital stopwatch. Data points are identified as individual’s values. Columns represent mean ± S.E.M. CTRL-CQ: uninfected and CQ- treated mice; INF-CQ: infected and CQ treated mice; FL1: familiar location 1; FL2: familiar location 2. Two-way RM ANOVA/Bonferroni was used for intragroup comparison of different objects at familiar locations (n=12). Representative of two experiments.

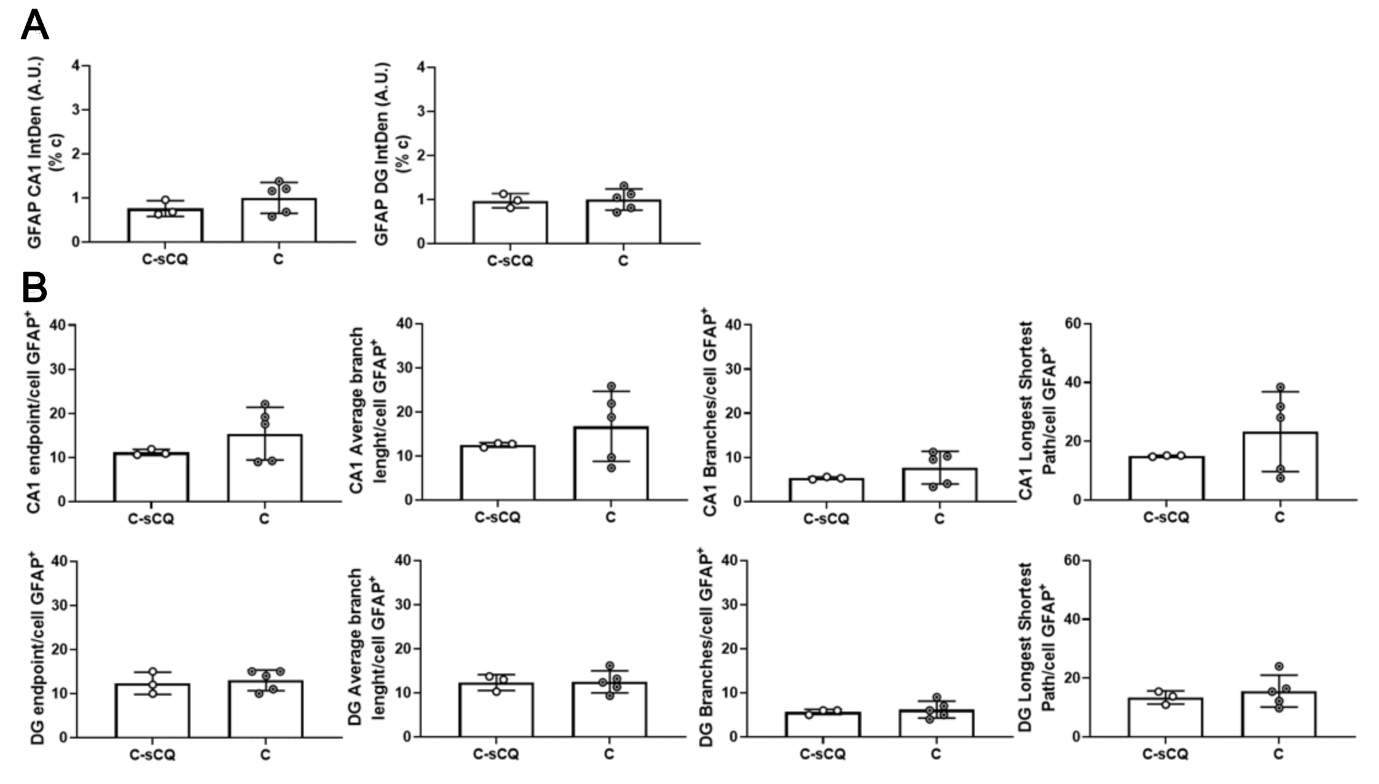

**Figure S2. Effect of chloroquine treatment on GFAP immunoreactivity of hippocampal cells and morphological analysis.** Immunoreactivity of GFAP cells was detected in dentate gyrus and CA1 of hippocampus of uninfected CQ treated (C) and uninfected PBS treated (C-sCQ) mice. (**A**) Quantification of integrated density of GFAP immunoreactivity of dentate gyrus and CA1. (**B**) Cytoskeleton analysis of GFAP^+^ cells in hippocampal dentate gyrus and CA1 regions. Columns represent mean ± standard deviation of uninfected and treated with PBS control (C-sCQ) and uninfected and CQ treated (c) (n=3-5). Data points are identified as individuals’ values. Columns represent mean ± S.D. Mann-Whitney test.

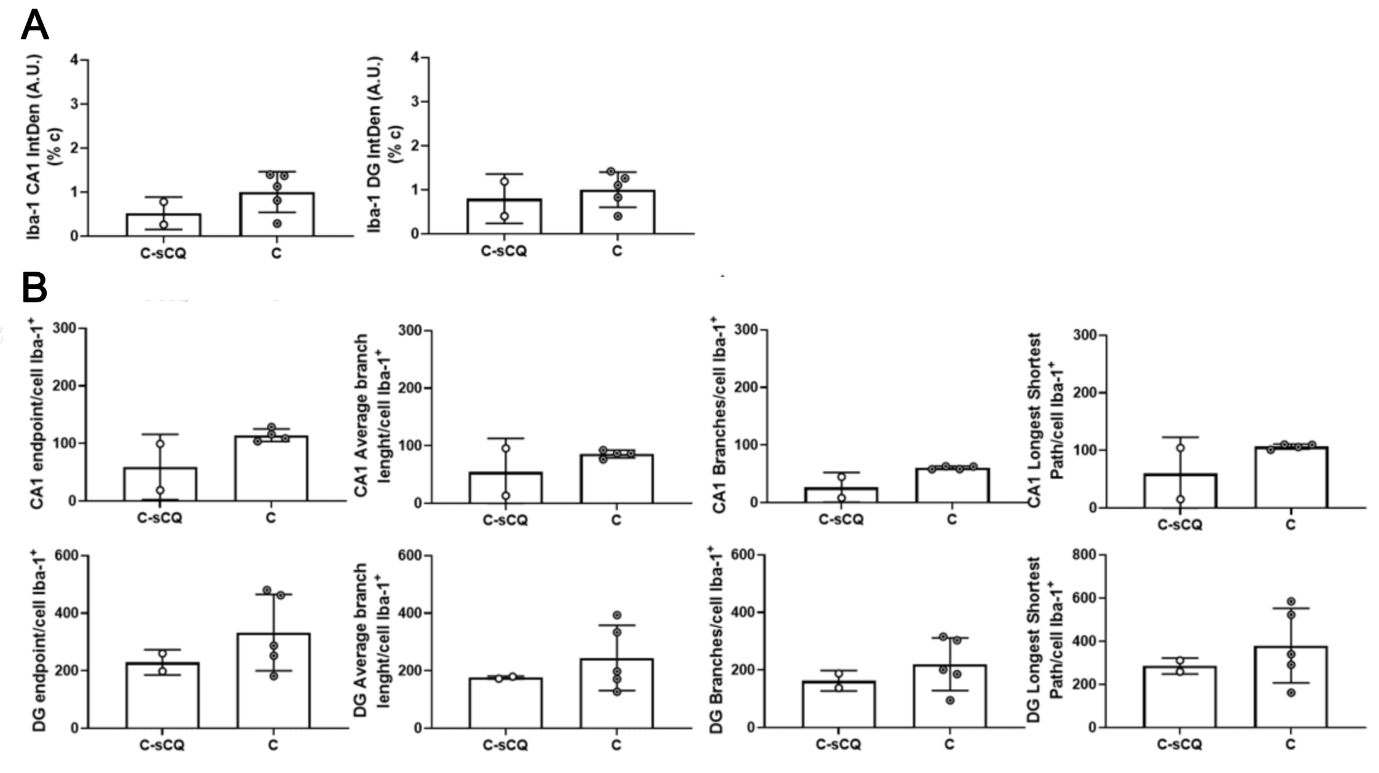

**Figure S3. Effect of chloroquine treatment on Iba-1 immunoreactivity of hippocampal cells and morphological analysis.** Immunoreactivity of Iba-1 cells was detected in dentate gyrus and CA1 of hippocampus of uninfected CQ treated (C) and uninfected PBS treated (C-sCQ) mice. (**A**) Quantification of integrated density of Iba-1 immunoreactivity of dentate gyrus and CA1. (**B**) Cytoskeleton analysis of Iba-1^+^ cells in hippocampal dentate gyrus and CA1 regions. Columns represent mean ± standard deviation of uninfected and treated with PBS control (C-sCQ) and uninfected and CQ treated (C) (n=2-5). Data points are identified as individuals’ values. Columns represent mean ± S.D. Mann-Whitney test.
